# Phenotype Discovery from Population Brain Imaging

**DOI:** 10.1101/2020.03.05.973172

**Authors:** Weikang Gong, Christian F. Beckmann, Stephen M. Smith

## Abstract

Neuroimaging allows for the non-invasive study of the brain in rich detail. Data-driven discovery of patterns of population variability in the brain has the potential to be extremely valuable for early disease diagnosis and understanding the brain. The resulting patterns can be used as imaging-derived phenotypes (IDPs), and may complement existing expert-curated IDPs. However, population datasets, comprising many different structural and functional imaging modalities from thousands of subjects, provide a computational challenge not previously addressed. Here, for the first time, a multimodal independent component analysis approach is presented that is scalable for data fusion of voxel-level neuroimaging data in the full UK Biobank (UKB) dataset, that will soon reach 100,000 imaged subjects. This new computational approach can estimate modes of population variability that enhance the ability to predict thousands of phenotypic and behavioural variables using data from UKB and the Human Connectome Project. A high-dimensional decomposition achieved improved predictive power compared with widely-used analysis strategies, single-modality decompositions and existing IDPs. In UKB data (14,503 subjects with 47 different data modalities), many interpretable associations with non-imaging phenotypes were identified, including multimodal spatial maps related to fluid intelligence, handedness and disease, in some cases where IDP-based approaches failed.

## 1 Introduction

Large-scale multimodal brain imaging has enormous potential for boosting epidemiological and neuroscientific studies, generating markers for early disease diagnosis and prediction of disease progression, and the understanding of human cognition, by means of linking to clinical or behavioural variables. Recent major studies have been acquiring brain magnetic resonance imaging (MRI), genetics and demographic/behavioural data from large cohorts. Examples are the UK Biobank (UKB)^1^, the Human Connectome Project (HCP)^2^ and the Adolescent Brain Cognitive Development (ABCD) study^3^. These studies involve multimodal data, meaning that several distinct types of MRI data are acquired, mapping activity, functional networks, structural connectivity, white matter microstructure, and organisation and volumes of different brain tissues and sub-structures^1^. However, the multimodal, high-dimensional and noisy nature of such big datasets makes many existing analytical approaches for extracting interpretable information impractical^4^.

Traditionally, large-scale neuroimaging studies first summarize the imaging data into interpretable image-derived phenotypes (IDPs)^1,5^, which are scalar quantities derived from raw imaging data (e.g., regional volumes from structural MRI, mean task activations from task MRI, resting-state functional connectivities between brain parcels). This knowledge-based approach is simple and efficient, and effectively reduces the high-dimensional data into interpretable, compact, convenient features. However, there may well be a large loss of information, due to such “expert-hand-designed” features not capturing important sources of subject variability (or even just losing sensitivity by the pre-defined spatial sub-areas being suboptimal), as well as ignoring cross-modality relationships. Further, such uni-modal compartmentalised analyses do not utilise the fact that for many biological effects of interest we expect there to be biological convergence across different data modalities, iė; changes in the underlying biological phenotype likely manifest themselves across multiple quantitative phenotypes, so that a joint analysis effectively increases both the power of detecting such effects and the interpretability of the findings.

In contrast to such uni-modal analyses, data-driven multivariate approaches (i.e., unsupervised machine learning) have been proposed, which perform simultaneous decomposition of voxel-level data directly, generally representing data as the summation of a number of “components” or “modes”. Each mode is formed as the outer product of two vectors: one is a vector of subject weights (describing the relative strength of expression of that mode in each subject), and a vector of voxel weights (in effect a spatial map for each data modality, describing the spatial localisation of the mode). The subject weight vectors (one per mode) can be considered “features”(similar to IDPs, but being data-driven) for use in further modelling, such as for the prediction of non-imaging variables. They are often either based on eigen-decomposition, such as multi-set canonical correlation analysis (mCCA)^6,7^, or based on variations of independent component analysis (ICA)^8–11^. Among them, FMRIB’s Linked ICA (FLICA)11 is an efficient approach which has been successfully applied to identify brain systems that are involved in lifespan development and diseases^12,13^, attention deficit hyperactivity disorder^14^, preterm brain development^15^ and cognition and psychopathology^16^. FLICA has advantages compared with uni-modal analysis on IDPs, including: (1) It leverages the cross-modality information of multimodal data, so has the potential to detect patterns that are not discoverable in any single modality; (2) It is a data-driven objective approach which automatically discovers meaningful patterns in voxel-level multimodal data by searching for spatial non-Gaussian sources that have been shown to likely reflect real structured features in neuroimaging data^17^. While this approach has been applied successfully to medium-sized cohort data^12–16^, the original algorithms for carrying out FLICA do not scale well with increasing data size, and are unable to analyze large datasets such as UKB, where dozens of different modalities over tens of thousands of subjects are available. Importantly, because the core FLICA algorithms are multivariate, acting in a complex way simultaneously across all subjects, modalities and voxels using Variational Bayesian updates of parameters, this problem cannot be solved through simple parallelisation or other algorithmically simple methods for distributing computations across a large cluster, and so cannot be addressed simply by increasing the number of processors or memory available.

To tackle this problem, we propose an approach that embeds advanced data compression techniques across the different data dimensions into the FLICA approach. We use a multimodal extension of MELODIC’s Incremental Group Principal component analysis^18^ (mMIGP, applied across modalities) and online dictionary learning^19^ (DicL, applied within-modalities) to efficiently reduce the size of multimodal neuroimaging data. The reduced data are then characterised through FLICA in terms of underlying modality-specifc maps and subject loading vectors. Here we refer to this combination of techniques as Big-data FLICA, or BigFLICA for short). Two important advantages of the proposed approach are: (1) Preserving key information in original data but also reducing the effects of stochastic domain-specific noise; (2) Increasing the computational efficiency of the FLICA algorithm for extremely large population datasets. BigFLICA is scalable for simultaneously analyzing all the multimodal data of the full 100,000-subjects UKB dataset using only a modest computing cluster (**Fig. 1**).

**Figure 1:**
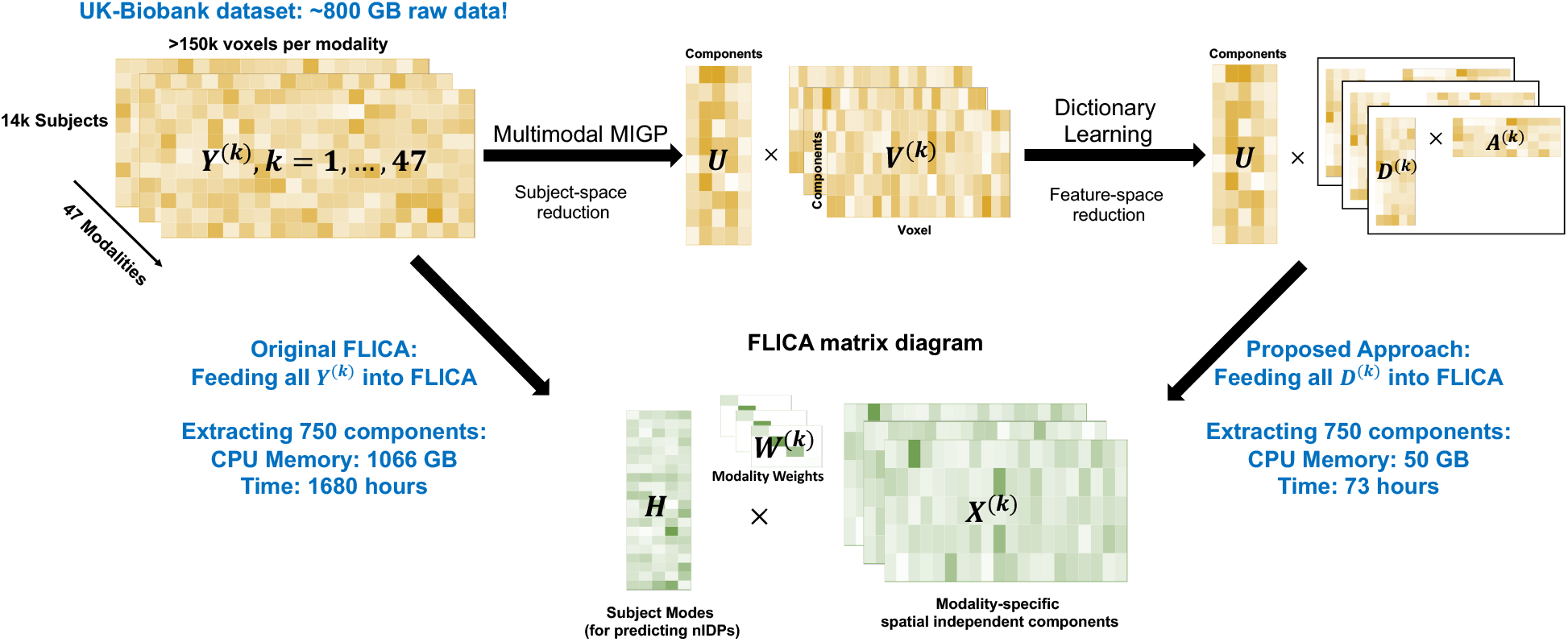
Overview of the proposed approach for jointly analyzing a biobank-scale multimodal neuroimaging dataset. Currently for the UKB dataset (voxel-level data, 14503 subjects, 47 modalities), the total data size is approximately 800 GB, and if we directly feed these data into FLICA and extract 750 components, we will need approximately 1066 GB CPU memory and 1680 hours computation time. Our new approach, BigFLICA, used multimodal MIGP and dictionary learning to preprocess the multimodal data; this is efficient and memory friendly, and much of this preprocessing can be easily parallelized. BigFLICA only used 50 GB memory and 73 hours to analyze the same dataset using a 24-core compute server.

We first demonstrate the effectiveness of our approach through extensive simulations. Then, in real data, we quantify performance in terms of the prediction accuracy of non-imaging-derived phenotypes (nIDPs)^20,21^, such as health outcome measures. Using voxel-level imaging data of 81 modalities from 1,003 subjects in the HCP and 47 modalities from 14,053 subjects in the UKB, we show that BigFLICA can perform comparably with original FLICA^11^ in terms of the prediction accuracy for nIDPs (158 in HCP and 8,787 in UKB). Most importantly, we systematically investigated whether there are benefits to jointly fusing multimodal data together, instead of analysing them separately. We show that significant improvements in the prediction accuracy of nIDPs are found when comparing a high-dimensional BigFLICA with other widely-used data analysis strategies: (1) doing single-modality ICA and concatenating the results across modalities and (2) using existing IDPs (5,812 in HCP and 3,913 in UKB). In particular, the improvements in prediction of many health outcome and cognitive variables are large, more than doubling prediction accuracy for some variables. Furthermore, we investigate the relationship between modes derived by BigFLICA and IDPs. We find that although the modes were estimated from the same set of voxel-level data, they have complementary information which can be combined together to further increase the prediction accuracy of nIDPs. Finally, we applied BigFLICA to analyze the UKB data and extracted 750 components. Existing multimodal ICA cannot estimate this many modes from this many subjects. We found several interpretable associations between modes of BigFLICA and nIDPs, including modes that relate to *fluid intelligence, handedness, age started wearing glasses or contact lenses* and *hypertension*. In many cases BigFLICA can find associations with nIDPs with greater statistical sensitivity than was possible with IDPs. Overall, BigFLICA demonstrated the advantages of data-driven joint multimodal modelling in the analysis of biobank-scale multimodal datasets.

## 2 Results

### Brief overview of the proposed approach: BigFLICA

FLICA^11^ is a Bayesian ICA approach for multimodal data fusion. The input of FLICA is *K* modalities’ data matrices *Y* ^(*k*)^ with dimensions *N* × *P_k_,k* = 1,…, *K*, where *P_k_* is the number of features (e.g., voxels) and *N* is the number of subjects. FLICA aims to find a joint *L*-dimensional decomposition of all *Y*^(*k*)^: *Y*^(*k*)^ = *HW*^(*k*)^*X* ^(*k*)^ + *E*^(*k*)^, where *H*_(*N*×*L*)_ is the shared subject mode (mixing matrix) across modalities (a vector of subject weights for each mode), so is a ‘link’ across different modalities, 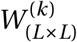 is a positive diagonal mode-weights matrix (one overall weight per modality per mode), 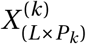 is the independent (spatial) feature maps for the *L* components of a modality (one map per modality per mode), and 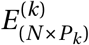 is the modality-specific Gaussian noise term (**Fig. 1**). We propose two efficient approaches that can either be used separately or combined together to reduce the size of the original data matrices, and therefore reduce the computational load of the original FLICA. An overview of BigFLICA is shown in **Fig. 1**.

The first approach, termed multimodal extension of MELODIC’s Incremental Group Principal component analysis^18^ (mMIGP), aims to reduce the subject dimension to a linear combination of the original subjects. mMIGP is a time- and memory-efficient approximation of principal component analysis (PCA) on feature-concatenated multimodal data. To this end, if we aim to get a *L*^★^ decomposition, we first apply MIGP^18^ separately within each modality to estimate 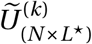, which is an approximation of an *L*^★^-dimensional PCA decomposition of one modality *Y*^(*k*)^. This step can be done in parallel across modalities. Then, we concatenate all 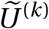 in the component dimension and apply another MIGP to get *U*_(*N*×*L*^★^)_, which is an *L*^★^-dimensional approximate PCA decomposition of all modalities together. Finally, we project the original data of each modality *Y*^(*k*)^ to the PCA-reduced space using*U*. If no further reduction (e.g., dictionary learning as below) is to be applied, the data that could then be fed into the core FLICA would be the *K* component-by-feature matrices *V*^(*k*)^ of size *L*^★^ × *P_k_*, and FLICA would then extract *L* (*L* < *L*^★^) components from these (**Methods**). This step almost adds little computational cost compared with the original FLICA, because a similar PCA step is needed to initialize the parameters of the original FLICA, but this approach is feasible for large numbers of subjects and modalities. Although different modalities usually have different overall signal-to-noise ratios (SNR), which is largely ignored by this mMIGP step, the subsequent FLICA can take this into account by the modality-specific noise terms, and a high-dimensional mMIGP is used to capture modes with even small variations in each modality.

It is known that voxels are correlated in both a local fashion (local spatial autocorrelation) and across brain networks (long range correlation); hence, effective feature subsampling could hope to capture all important information in the data but also reduce the cost of spatial modelling in FLICA^22^. Therefore, we incorporate an approach, termed sparse online Dictionary Learning^19^ (DicL), to reduce the dimension of feature (e.g., voxel) space that can capture both local and distant spatial correlation structure. Specifically, for each modality, we use DicL to model the *V*^(*k*)^ as a sparse linear combination of *L*^★★^ basis elements: *V*^(*k*)^ = *A*^(*k*)^*D*^(*k*)^, where *D*^(*k*)^ is the *sparse spatial dictionary basis*, and *A*^(*k*)^ is the *feature loadings*. By minimizing the reconstruction error, and enforcing sparsity in the dictionary basis *D*^(*k*)^, we aim to achieve an optimal subsampling of feature space. The inputs of FLICA are then *K* smaller matrices *A*^(*k*)^, which are only of dimension *L*^★^ × *L*^★★^, and FLICA then extracts *L* (*L* < min(*L*^★^, *L*^★★^)) components from these (**Methods**). Compared with doing FLICA with the original *K* large *N* × *P_k_* matrices, using the DicL preprocessed data can greatly reduce the computation load of FLICA. DicL can easily be parallelized across modalities and is memory friendly, which further increases efficiency (**Fig. 1**).

### Evaluation of BigFLICA in simulations

We first applied BigFLICA on simulated data to evaluate the performance of mMIGP and DicL as data preprocessing approaches under different parameter settings and data signal-to-noise ratios. The mean correlation of extracted components with simulated ground truth was compared with the corresponding result from the original FLICA (**Methods**).

For mMIGP, **Fig. 2a** shows that, in most of the situations, the BigFLICA with mMIGP preprocessing gave similar results to the original FLICA, and both FLICA and BigFLICA accurately find the underlying ground truth in most cases. This is in agreement with results of simulations in the MIGP paper^18^ that it can accurately approximate a full-data PCA in different situations. The optimal dimension of mMIGP is different among simulations; sometimes a relative low dimension can achieve an accurate estimation of components (e.g. **Fig. 2a** first three columns), while in other cases a high dimension is needed (e.g. **Fig. 2a** the fourth column).

**Figure 2:**
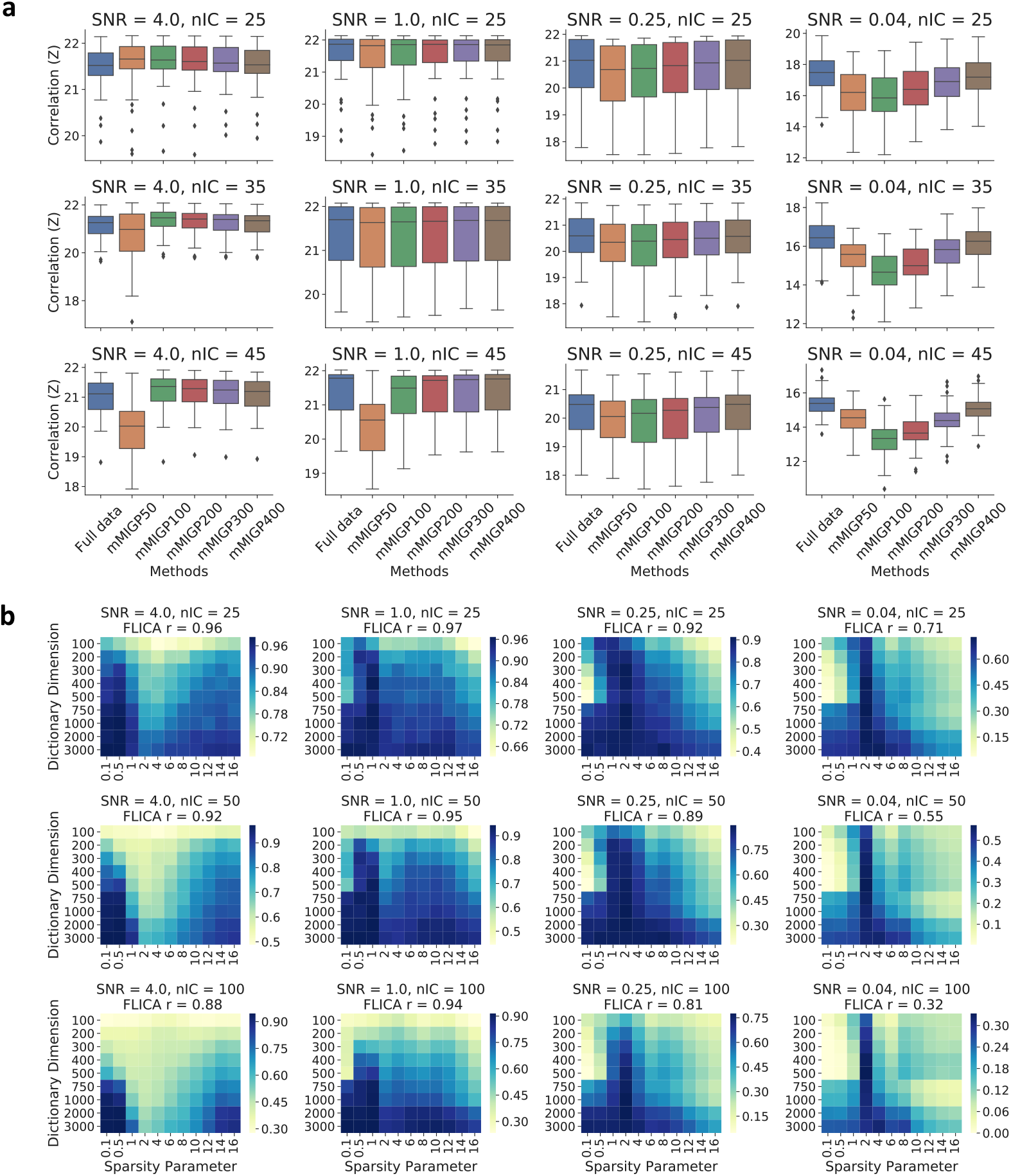
Evaluation of multimodal extension of MIGP (mMIGP) and dictionary learning (DicL) as the data preprocessing steps for the FLICA using simulations. BigFLICA achieves similar performance as compared with original FLICA that uses the full data. a, Evaluation of mMIGP preprocessing. We compared the correlations (Z-transformed) of extracted components with ground truth across 50 simulations using the original FLICA (the left column of each figure) and the mMIGP preprocessed FLICA (other columns). The mMIGP dimensions vary between 50 and 400; the SNRs are between 4 and 0.04 (left to right), and the number of FLICA and ground truth components are 25, 35, 45 (top to bottom). As there are 500 subjects, the reduction factor is from 10 to 1.25. b, Evaluation of DicL preprocessing. We compared the the correlations of extracted components with ground truth using the original FLICA (FLICA results given in the titles of each figure) and the DicL preprocessed FLICA with different sparsity parameters and dictionary dimensions (cells of the heatmaps). The SNRs are between 4 and 0.04 (left to right), and the number of FLICA and ground truth components are 25, 50, 100 (top to bottom). As there are 27,000 original features per modality, the reduction factor is from 270 to 9.

For DicL, **Fig. 2b** shows that in almost all circumstances: (1) increasing the dictionary dimensions will boost the performance of subsequent FLIC Aanalysis; (2) the optimal sparsity parameters are usually between *λ* = 0.5 to 2, and they have similar performance; (3) In most cases the optimal performance given by DicL matches that of non-reduced analysis (noted in figure legends). Therefore, in the real data analysis, when using the DicL approach, we always use a very high dimensional DicL decomposition and fix the sparsity parameter to *λ* = 1.

### Computation time comparison

**Table 1** shows the comparison of the computation time and memory requirement of BigFLICA with the original FLICA in the UKB dataset. All code was implemented in Python 2.7, and both BigFLICA and FLICA were run using 24 cores on a single compute node with Intel Xeon CPU E7-8857 v2 @ 3.00GHz CPU and 2048 GB RAM. The computation time includes: (1) Preprocessing of data using mMIGP and DicL (BigFLICA only); (2) Initialization of FLICA parameters; (3) FLICA VB parameter updates. For the 100,000-subjects data, BigFLICA greatly decreases the computation time and memory usage from an unrealistic amount to a modest configuration for a modern HPC cluster, which therefore allows for the possibility of data-driven population phenotype discovery.

**Table 1:**
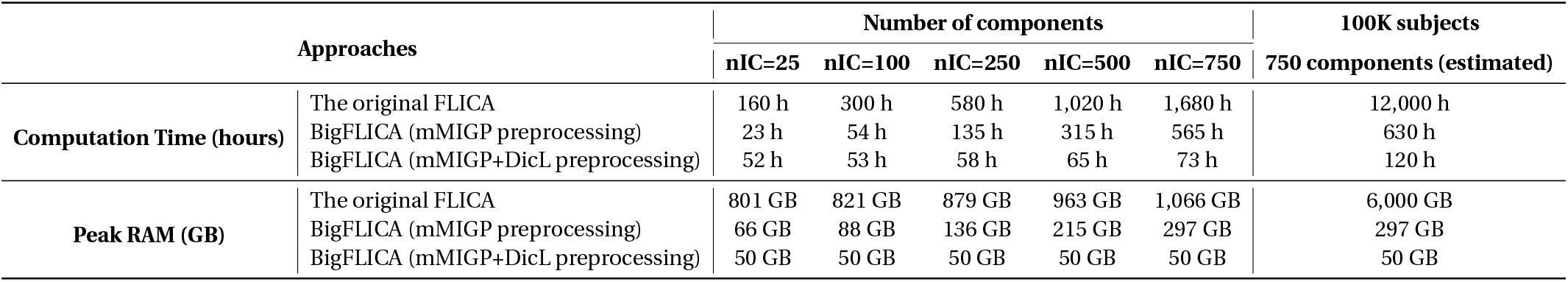
Comparison of computation time and amount of RAM usage of BigFLICA with the original FLICA in the UKB dataset (14,503 subjects, 47 different modalities). BigFLICA greatly increases computational efficiency in different settings. Both BigFLICA and FLICA were run on the same computer using all 24 cores in all computation stages with Intel Xeon CPU E7-8857 v2 @ 3.00GHz and 2TB RAM.

| Approaches |  | Number of components |  |  |  |  | 100K subjects |
| --- | --- | --- | --- | --- | --- | --- | --- |
|  |  | nIC=25 | nIC=100 | nIC=250 | nIC=500 | nIC=750 | 750 components (estimated) |
| Computation Time (hours) | The original FLICA | 160 h | 300 h | 580 h | 1,020 h | 1,680 h | 12,000 h |
|  | BigFLICA (mMIGP preprocessing) | 23 h | 54 h | 135 h | 315 h | 565 h | 630 h |
|  | BigFLICA (mMIGP+DicL preprocessing) | 52 h | 53 h | 58 h | 65 h | 73 h | 120 h |
| Peak RAM (GB) | The original FLICA | 801 GB | 821 GB | 879 GB | 963 GB | 1,066 GB | 6,000 GB |
|  | BigFLICA (mMIGP preprocessing) | 66 GB | 88 GB | 136 GB | 215 GB | 297 GB | 297 GB |
|  | BigFLICA (mMIGP+DicL preprocessing) | 50 GB | 50 GB | 50 GB | 50 GB | 50 GB | 50 GB |

### Real data: Comparing BigFLICA with the original FLICA based on the prediction accuracy of nIDPs

As there is no ground truth available, we tested modes of BigFLICA have a similar prediction accuracy of nIDPs compared with the original FLICA, using data from the HCP, and a subset of 1,036 subjects from the UKB. Elastic-net regression with nested 5-fold cross-validation was used to predict each of the nIDPs. This approach is widely-used and has been shown to achieve a robust and state-of-the-art performance in many neuroimaging studies^24,25^. Pearson correlation between each of the predicted and the true nIDPs in the outer test fold is used to quantify accuracy. The statistical significance of differences of prediction accuracy between two approaches are estimated by a weighted paired t-test approach. (**Methods**).

**Fig. 3** shows the Bland-Altman plots comparing the prediction accuracy of nIDPs between original FLICA and BigFLICA with mMIGP preprocessing only (**Fig. 3a**), and with DicL preprocessing only (**Fig. 3b**), and with both data reduction approaches (**Fig. 3c**), in the UKB and HCP datasets. In these comparisons, mMIGP reduced the data to approximately 1/10 to 1/2 of the original data size, and DicL reduced data to approximately 1/75 of the original data size. Overall, BigFLICA can estimate similar sets of modes with comparable prediction accuracy in real multimodal neuroimaging data, i.e., the difference of the correlation between two methods is centered around zero across a wide range of mean correlation values (which are also reflected in the insignificant p-values of weighted paired t-test), which demonstrates that the mMIGP and DicL approaches are effective to reduce data and preserve key information in the data.

**Figure 3:**
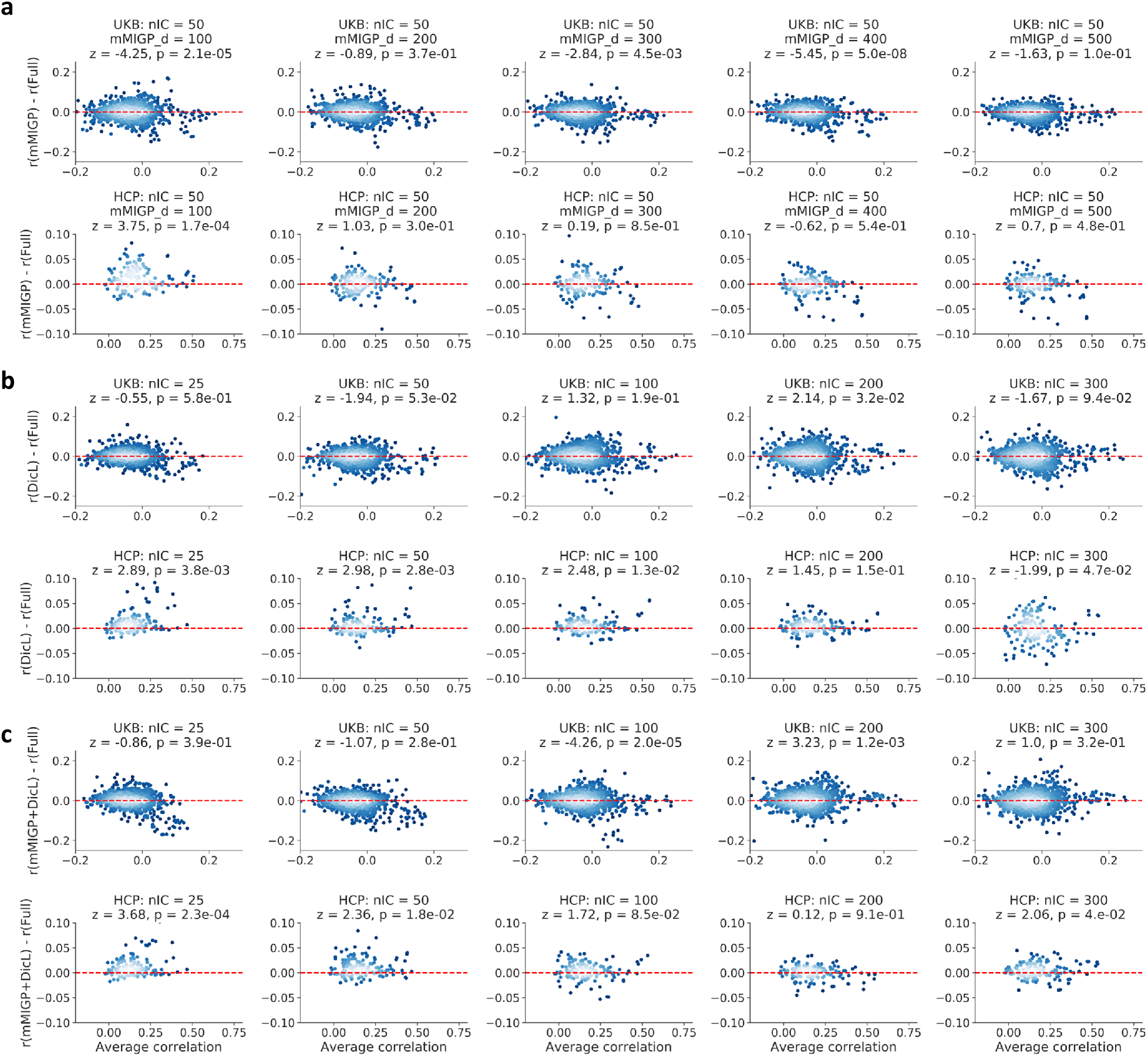
Comparison of prediction accuracy of nIDPs between BigFLICA and the original FLICA. Overall, for most of the comparisons, the differences of prediction accuracy are not significant. In each of the Bland–Altman plots, each point represents the prediction of one nIDP, where the x-axis is the average prediction correlation of the two approaches while the y-axis is the difference, i.e., BigFLICA - FLICA. The z- and p-values in the titles reflected the statistical significance of the differences. The Bonferroni correction 0.05 threshold corresponds to a raw p-value of 1.7e-3. a, Comparing FLICA with mMIGP preprocessing with the original FLICA. We used a subset of 1,036 subjects in the UKB dataset (top) and the HCP (bottom). The number of estimated FLICA components is set to 50, and mMIGP dimensions are set from 100 to 500. b, Comparing FLICA with DicL preprocessing with the original FLICA. We used a subset of 1,036 subjects in the UKB dataset (top) and the HCP (bottom). The dictionary dimension is set to a high value of 2000, and the sparsity parameter is set to *λ* = 1 for all modalities. The number of estimated FLICA components are set from 25 to 300. c, Comparing FLICA with both mMIGP and DicL preprocessing combined, with the original FLICA. The mMIGP dimension is set to 500, and other settings are the same as in b. We use only a subset of UKB here so that running the original FLICA is computationally feasible.

### Comparing BigFLICA with multiple independent single-modality ICA decomposition

We also compared BigFLICA outputs against features pooled across those from separate ICA processing of each modality. **Fig. 4a** shows that BigFLICA has a worse prediction performance than via running ICA separately on each modality when the dimensionality *L* is low. This is because at low dimensional decomposition, single-modality ICA is most efficient because the constraints imposed on the degrees-of freedom implied in the FLICA model is insufficient to capture the important data variation into joint components. However, when *L* becomes large, the prediction accuracy becomes better than the single-modality ICA (e.g., ⩾ 250 in UKB). This is because, at high dimensional decomposition, BigFLICA effectively combines multimodal information by considering cross-modal correlation in the data decomposition stage. Although the cross-modal correlation is considered in the final prediction stage when using single-modality ICA, the fact that BigFLICA identifies and takes advantage of correlated information between modalities at an earlier stage in feature generation helps improve the prediction performance.

**Figure 4:**
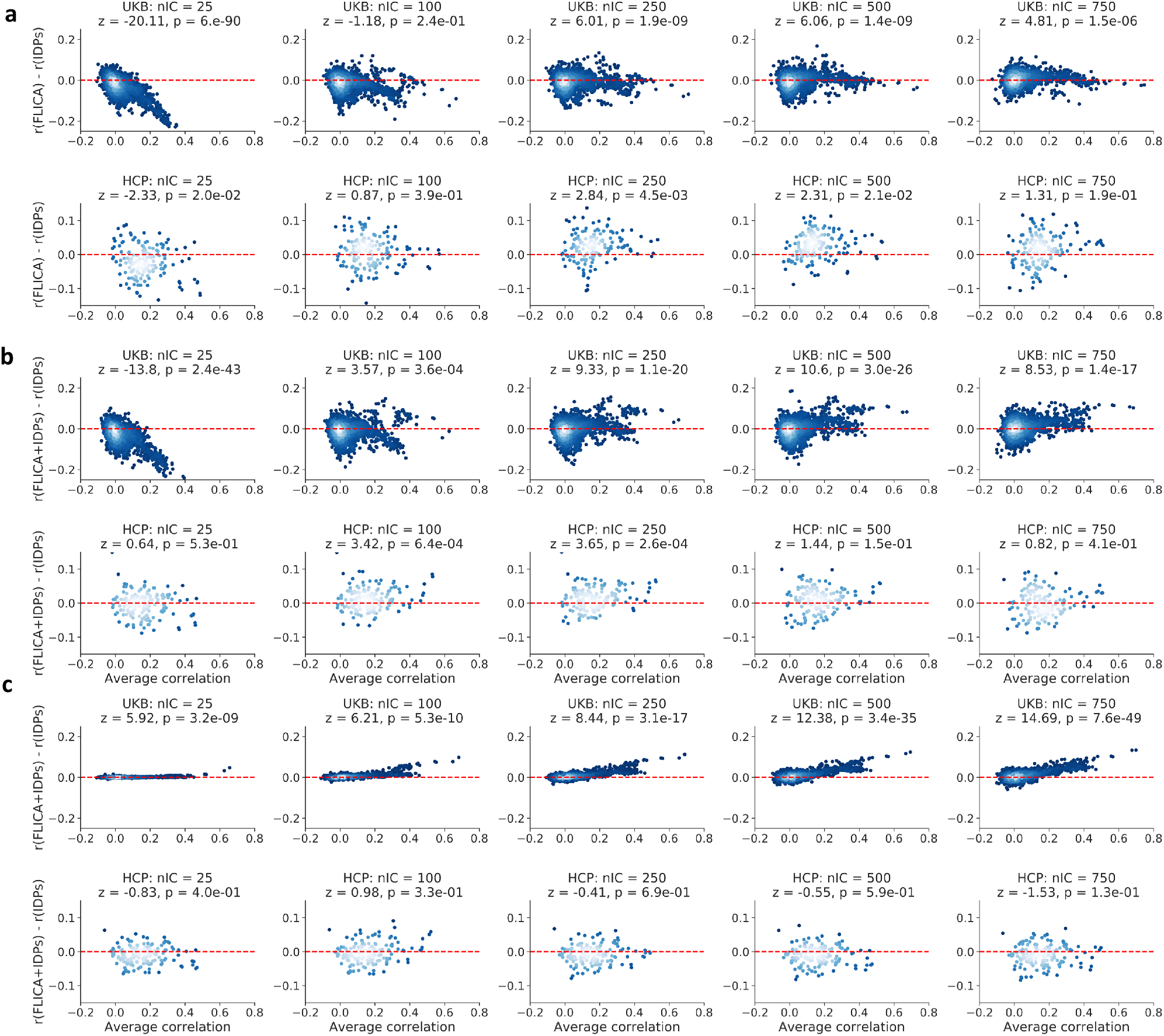
Comparison of prediction accuracy of nIDPs between BigFLICA against single-modality ICA and the IDPs. Overall, for high-dimensional BigFLICA decompositions in the UKB dataset, BigFLICA achieved statistical significant increases of prediction accuracy of nIDPs compared with single-modality ICA and IDPs. Combining BigFLICA and IDPs together future improves compared with IDPs alone. In each of the Bland–Altman plots, each point represents the prediction of an nIDP, where the x-axis is the average prediction correlation of the two approaches, while the y-axis is the difference. The z- and p-values in the titles reflected the statistical significance of the differences. The Bonferroni correction 0.05 threshold corresponds to a raw p-value of 1.7e-3. a, Comparing BigFLICA with the concatenation of single-modality ICA outputs. Top: UKB; Bottom: HCP. The number of FLICA components is set from 25 to 750. b, Comparing BigFLICA with IDPs. Top: UKB; Bottom: HCP. The number of IDPs is 3,913 in UKB and 5,812 in the HCP. c, Comparing the concatenation of BigFLICA and IDPs against IDPs only. Top: UKB; Bottom: HCP.

### Comparing BigFLICA with hand-curated imaging-derive phenotypes

We compared the predictive performance of BigFLICA with IDPs in both HCP and UKB datasets (**Methods**). **Fig. 4b** shows that, in the UKB data, when the number of modes is low, BigFLICA has a worse predictive power than the joint performance of 3,913 IDPs, due to the same insufficient degree-of-freedom reason as above. However, when the dimensionality becomes higher, BigFLICA is clearly outperforming the IDPs, owing to jointly fusing multimodal voxelwise data by considering cross-modality correlation. In the HCP data, the performance is overall similar. These results indicate that BigFLICA can potentially explain more pheno-typic and behavioural variances than IDPs.

In more detail, **Tables S1** shows that, in the UKB dataset, the high-dimensional BigFLICA (nIC=750) has improved prediction accuracy for many nIDPs that relate to *cognition phenotypes* and *health outcomes* compared with IDPs. These tables do not include nIDPs where both methods have low predictive power (*r* < 0.1). In the HCP dataset (**Table S2**), BigFLICA (nIC=100) also shows improved prediction accuracy in many cognitive and health outcomes variables compared with using IDPs.

Further, when we concatenated the modes of BigFLICA and IDPs together to predict nIDPs, as shown in **Fig. 4c**, the combined feature sets have a significant improvement of prediction accuracy than the IDPs alone in the UKB data. There are almost no differences for the same comparison in the HCP data. This suggests that BigFLICA and IDPs may contain some complementary information of nIDPs.

To investigate the relationships between BigFLICA and IDPs further, we built prediction models that used modes of BigFLICA to predict each of the IDPs, to further characterise information overlap and complementarity between the two approaches. As shown in **Figs. S2a** and **S2b**, different types of IDPs can be predicted differently, and the resting-state functional connectivities always had the worst accuracy in both the HCP and the UKB datasets, because they are (relatively) noisy. However, when using BigFLICA modes to predict 6 new summary features of the connectivity matrices (derived by applying ICA to the matrix of subjects by network matrix edges)^5^, the accuracy is very high (*r* range from 0.85 to 0.89 for a 100 dimensional BigFLICA decomposition). In addition, when we used IDPs to predict modes of BigFLICA, as shown in **Figs. S2c** and **S2d**, the prediction correlation almost showed a bimodal distribution, which means that some of the FLICA modes can be predicted by the IDPs (mean *r* ≈ 0.8) while others cannot (mean *r* ≈ 0.2). These results further demonstrates that BigFLICA and IDPs span significant complementary variance.

### BigFLICA comparison with mCCA and reproducibility

We next compared BigFLICA against mCCA (eigendecomposition based modelling, which of course also would require similar advances to BigFLICA in order to work on large data; see online Methods). Overall, BigFLICA had (very slightly) improved prediction accuracy (**Fig. S3**), and with slightly more parsimonious modelling (**Fig. S4**). However, with split-half (across subjects) reproducibility testing, BigFLICA components were considerably more reproducible than those from mCCA (median BigFLICA correlation greater than 0.9 in all cases, while many mCCA dimensionalities have median correlation less than 0.5) (**Fig. S5**).

### Examples of BigFLICA modes in the 14k UKB dataset

We now give four examples of significant associations between BigFLICA modes and nIDPs, namely, *Fluid intelligence*, *Age started wearing glasses or contact lenses, Handedness* and *hypertension*. In **Fig. 5**, we show the top four most strongly associated modalities in FLICA modes that correlate with a given nIDP. **Fig. S6** shows the population cross-subject mean maps for several task and rest fMRI modalities fed into FLICA. This helps give interpretive context for the FLICA mode maps, which depict subject variability in the activity/connectivity relative to these group mean maps.

**Figure 5:**
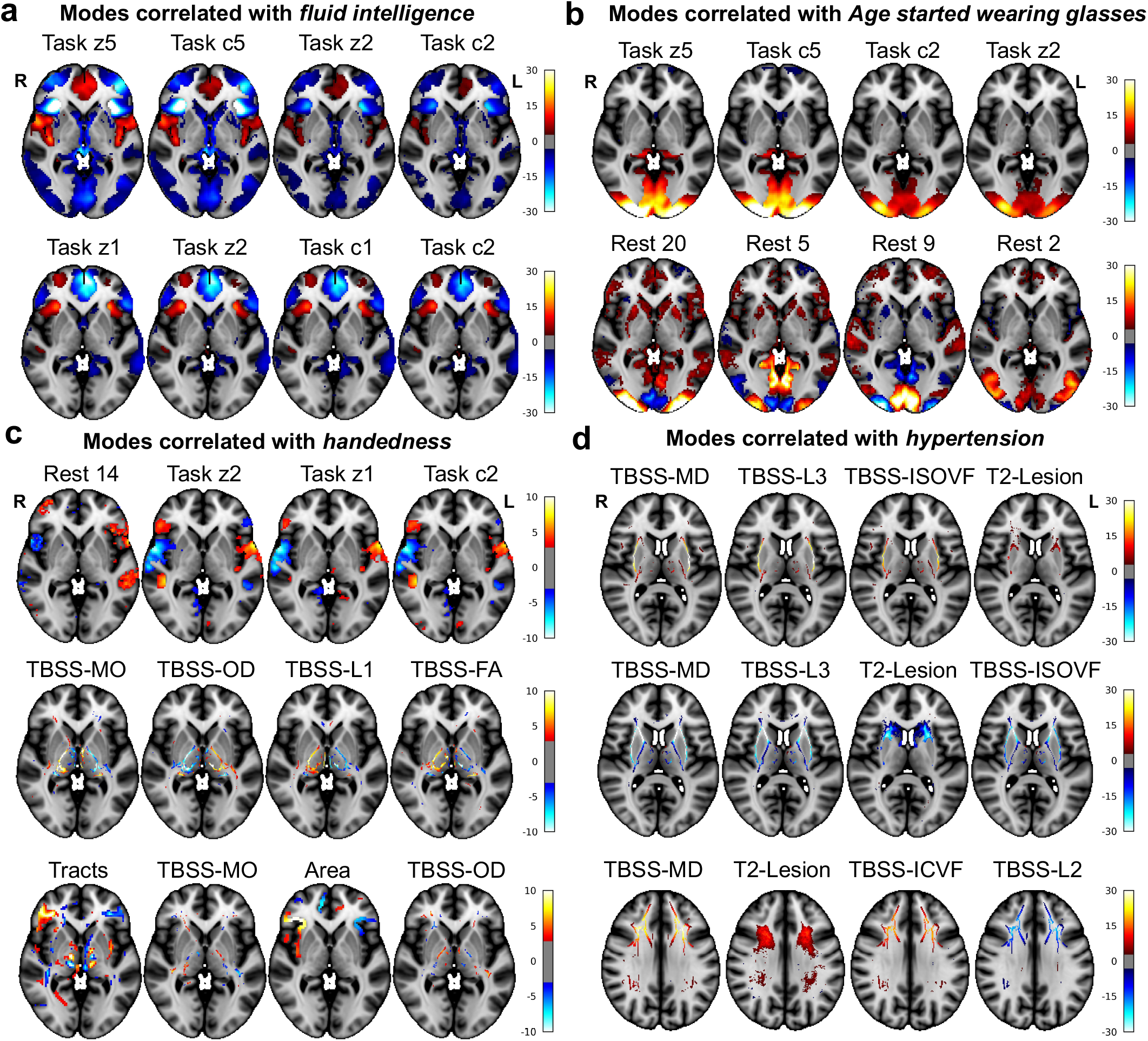
Examples of BigFLICA modes in the 14k UKB dataset. For each subfigure, each row shows one IC (BigFLICA mode or independent component) with top 4 most strongly associated modalities. **a,** Two BigFLICA modes that significantly correlate with *fluid intelligence* (IC25: *r* = −0.14; IC57: *r* = −0.12). **b,** Two BigFLICA modes that significantly correlate with *Age started wearing glasses or contact lenses* (IC164: *r* = −0.10; IC13: *r* = −0.05). **c,** Three BigFLICA modes that significantly correlate with *handedness* (IC235: *r* = −0.23; IC569: *r* = 0.07; IC232: *r* = −0.04). **d,** Three BigFLICA modes that significantly correlate with *hypertension* (IC259: *r* = 0.12; IC13: *r* = −0.11; IC319: *r* = −0.09). The Bonferroni corrected 0.05 threshold corresponds to an uncorrected p-value of 9.2 ×10^-9^ (corrected for number of components (750) and number of nIDPs (7245)). All of the above correlations passed the Bonferroni threshold except for IC232 with uncorrected *p* = 2.1 × 10^-7^.

For *Fluid intelligence*, using all modes (ICs) from the 750 dimensional BigFLICA decomposition as features (predictors) in multivariate elastic net prediction, a cross-validated prediction correlation of *r* = 0.26 is achieved. When we correlated each of the BigFLICA modes and IDPs with the fluid intelligence score in the UKB, we found that several task-fMRI-related BigFLICA modes have the strongest associations (**Fig. 5a**). The first (IC 25) involves task contrast “faces” and “faces>shapes” and the second (IC 57) involves contrast “shapes” and “face” (see **Tables S5** for the full list of these modalities). As the correlation of the mode IC 25 (i.e., its subject weights vector) with fluid intelligence is negative (r=-0.14), this means that the negative-weights voxels (such as in the anterior insula) are positively correlated with intelligence. The fMRI task (Hariri faces-shapes matching^29^) has, as expected, the greatest population average activation in sensory-motor areas (plus some amygdala involvement due to the emotionally negative nature of the faces), as seen in **Fig. S6**. However, the main brain areas involved in these modes are distinct, including anterior cingulate cortex, frontal pole, inferior frontal gyrus, and anterior insula; it is therefore interesting that the areas found by BigFLICA to be modulated in these components (and found to associate with intelligence) are more “frontal, cognitive” areas than the sensory-motor areas primarily activated on average. The top associations between fluid intelligence and IDPs also involve task-fMRI IDPs (**Tables S3**), but these were a factor of two weaker than associations with BigFLICA modes.

For *Age started wearing glasses or contact lenses*, BigFLICA achieved a prediction correlation of *r* = 0.16. Several resting-state connectivty and task modalities showed associations in primary visual areas (**Fig. 5b**), which is consistent with the fact that this is a vision-related health variable. Lower age of first wearing glasses is correlated with stronger activity in primary visual areas, and also with strength of resting-fMRI connectivity (or functional coherence) within the relevant areas of group-average connectivity; interestingly, in nearby distinct (but still primary visual) areas, there is reduction of correlation (blue voxels), suggesting greater differentiation of primary visual areas.

For *Handedness*, BigFLICA achieved a prediction correlation of *r* =0.23. BigFLICA identified several multimodal, lateralized (or laterally asymmetric) modes, including resting-state mode 14 (left-lateralised language network), task, surface area and white matter tracts (**Fig. 5c**). There are several resting-state connectivity-related IDPs correlated with handedness (**Tables S3**), consistent with a recent study^30^ that also used UKB IDPs, while no IDPs related to other modalities are found significant; in both cases the maximum IDP correlation only reached r=0.12, whereas the strongest association with BigFLICA modes was almost double this.

For a health variable *hypertension* (**Fig. 5d**), BigFLICA achieved a prediction correlation of *r* = 0.22. Several TBSS-related modalities showed consistent associations in the External Capsule tracts. Meanwhile, white matter hyperintensity (T2-Lesion volume) in the corresponding areas is also higher in people with hypertension. Several consistent findings have been reported in the literature^31–33^.

## 3 Discussion

In this paper, we presented BigFLICA, a multimodal data fusion approach which is scalable and tuneable to analyze the full UK-Biobank neuroimaging dataset, and other large-scale multimodal imaging studies. To the best of our knowledge, this is the first approach for data-driven (unsupervised) multimodal analysis in a brain imaging dataset of this size and complexity. Building on the top of the powerful FLICA model, we proposed a two-stage dimension-reduction approach that combines an incremental group-PCA (mMIGP) and dictionary learning (DicL) to effectively preprocess the multimodal dataset and reduce the computational load of the final FLICA, while maintaining or even improving performance, with as much as a 150-fold “intelligent” reduction in data size. We provide effective ways of choosing the hyper-parameters of BigFLICA, so that it is free of tuning except for choosing the final number of estimated components. Although this approach is motivated by the need for analyzing extremely big neuroimaging data, it is also applicable to other kinds of data such as genetics and behavioural measures. An easy-to-use version of this software will be integrated into an upcoming version of the FSL software package^34,35^. BigFLICA results on UKB will also be released via the UKB database as new data-driven IDPs (image features), for further epidemiological and neuroscientific research.

A strength of our work is that, unlike previous work that was limited to more moderate datasets and a few phenotypic and behavioural variables^8–11,36^, we used two of the largest, high-quality multimodal neuroimaging datasets, and thousands of phenotypic and behavioural variables to validate the proposed approach. We demonstrated that BigFLICA is not only much faster than the original FLICA (and can be run on very large data that is simply not analysable with FLICA or other existing methods), but also estimates similar modes with a comparable performance for predicting the non-imaging-derived phenotypes in real data (when tested on a large data subset that is just small enough to allow for comparison against FLICA). We provide insights into the advantages of data-driven multimodal fusion in big datasets by quantitative analysis^37,38^. First, when comparing BigFLICA with simpler IDP-based approaches (and also single-modality ICA approaches), we demonstrated that a high-dimensional BigFLICA has improved predictive power overall. We demonstrated the value of multimodal fusion instead of analyzing each modality separately. Second, when combining high-dimensional BigFLICA-derived features with IDPs together, the predictive power increased further compared with using either method alone. In addition, when we used BigFLICA-derived features and manually created (with expert knowledge) IDPs to predict each other, they cannot predict each other perfectly (although they are derived from the same imaging data). This indicates that BigFLICA-derived features and IDPs can be complementary to eacho ther, both therefore providing potentially important imaging biomarkers that capture different signal in the imaging data. An interesting finding is that although a high-dimensional BigFLICA has a much higher predictive power than a low dimensional decomposition, a low dimensional decomposition can still explain more than 80% of the total variance of the high dimensional decomposition. This suggests that some the phenotypic and behavioural variables are explained by only small proportions of variance of imaging data. Finally, in addition to the value of using BigFLICA-derived features for relating imaging to non-imaging data, BigFLICA components (particularly at lower dimensionalities) may allow us to learn more about how the different brain imaging modalities (and hence different spatial and biological aspects of the brain’s structure and function) relate to each other.

We see opportunities to improve the current approach. First, BigFLICA is limited to linear feature estimation, while the “ideal, true” information in imaging data may be highly nonlinear. Therefore, a nonlinear extension of BigFLICA, which might be achieved with kernel methods or deep neural networks, is an important area of further research. Second, BigFLICA is an unsupervised dimension reduction and feature generation approach. However, integrating some supervision, i.e., the target variable (such as disease outcomes), into the dimension reduction may boost the performance of the algorithm. Additionally, because BigFLICA generates data-driven features, as opposed to expert-created IDPs, the biological or anatomical interpretation of features is often likely not to be immediately obvious, requiring potentially intensive expert study. Future work could attempt to automate this interpretation process, for example by relating features to existing anatomical templates and atlases, and even by mining imaging literature. Finally, BigFLICA, or extensions, may be an effective way of discovering imaging confound factors^39^ that cannot be found by traditional approaches.

## Acknowledgments

We are grateful to UK Biobank and its participants (access application 8107). We thank the WU-Minn HCP Consortium for their invaluable contributions in generating the publicly available HCP data and implementing the procedures needed to acquire, analyze, visualize and share these datasets. Computation used the Oxford Biomedical Research Computing (BMRC) facility, a joint development between the Wellcome Centre for Human Genetics and the Big Data Institute supported by Health Data Research UK and the NIHR Oxford Biomedical Research Centre. Financial support was provided by the Wellcome Trust Core Award Grant Number 203141/Z/16/Z.

## Author contributions

W.G, C.B and S.S proposed the study, formulated the model, analyzed the data and wrote the manuscript.

## Competing interests

The authors declare that they have no competing financial interests.

## Data availability

BigFLICA will be released in an upcoming version of FSL. BigFLICA-derived features will be available from the UK Biobank database. For UK Biobank, all source data (including raw and processed brain imaging data, derived IDPs, and non-imaging measures) is available from UK Biobank via their standard data access procedure (see http://www.ukbiobank.ac.uk/register-apply). For HCP, data can be downloaded via website (http://humanconnectome.org/data) and ConnectomeDB. Matlab software for performing prediction using elastic-net regression is available at https://github.com/vidaurre/NetsPredict.

## Methods

### FLICA model

The input to FLICA is *K* modalities’ data matrices *Y*^(*k*)^ with each modality’s dimensions being *N* × *P_k_,k* = 1,…,*K*, where *P_k_* is the number of features (e.g., voxels) in modality *k* and *N* is the number of subjects. FLICA aims to find a joint *L*-dimensional decomposition of all *Y*^(*k*)^:

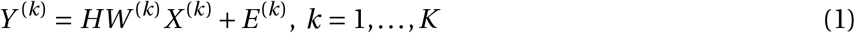

where *H*_(*N*×*L*)_ is the shared subject mode (mixing matrix) across modalities, so is a ‘link’ across different modalities, 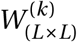 is a positive diagonal mode-weights matrix, 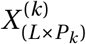 is the independent (spatial) feature maps forthe *L* components of modality *k*, and 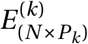 is the Gaussian noise term.

### Multimodal extension of MELODIC’s Incremental Group Principal component analysis for subject-space dimension reduction

We propose a multimodal extension of our previous MIGP approach^18^, termed mMIGP, to reduce the subject dimension of multimodal data. MIGP has been extensively validated in simulations and real neuroimaging data for finding an approximate PCA decomposition in a time- and memory-efficient way^18^. Suppose that our multimodal data are *K* matrices *Y*^(*k*)^,*k* =1,…,*K* with dimensions *N* × *P_k_*, where *N* is the number of subjects and *P_k_* is the number of features (e.g. voxels) in a modality. In mMIGP, each feature is z-score normalized first. Then, an MIGP is applied to each modality separately to find an *L*^★^-dimensional approximate PCA decomposition. Specifically, we want to find an approximation of a singular value decomposition (SVD) of each *Y*^(*k*)^:

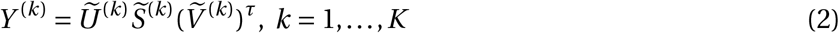

where 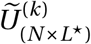 and 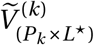 are the left and right singular vectors, while 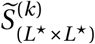 are the singular values. A naive SVD on *Y*^(*k*)^ scales quadratically with *N*, which is not efficient when *N* is large. To find the approximation, MIGP sequentially feed a subset of (columns of) *Y*^(*k*)^ in to an SVD, so that these subsets are reduced to a low-dimensional representation. The low-dimensional representation is then concatenated with another subset of (columns of) *Y*^(*k*)^, and is fed into another SVD to find the low-dimensional representation of them. The final SVD approximation is found after one pass of all data. The computational complexity of MIPG scales linearly with *N*. For a detailed description, please see Appendix A of the MIGP paper^18^.

The third step is to concatenate all 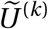 in the component dimension and apply another MIGP for finding a *L*^★^-dimensional approximate PCA decompositions *U* of size *N*×*L*^★^, which is a low-dimensional representation of multimodal data in the analysis. Finally, the z-score normalized data *Y*^(*k*)^ of each modality is projected onto the *U* by:

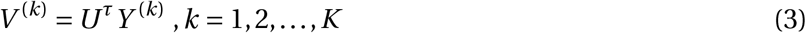

the *V*^(*k*)^, *k* = 1,2,…, *K* are the inputs of subsequent FLICA algorithm. Therefore, the total size of data output by this stage is 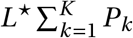, which is smaller than the original input size 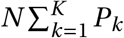. The fractional reduced data size is *L*^★^/*N*, and the *L*^★^ can be fixed when more subjects are introduced, so it is scalable in the big-data analysis. In practice, we usually choose *L*^★^ based on the percentage of explained variance of SVD in the third step.

If we feed *V*^(*k*)^, *k* = 1,2,…, *K* into FLICA to estimate *L*^★^ FLICA modes, the output subject mode matrix *H*^★^ is of the size *L*^★^ × *L*, so we then simply multiply this by *U* to get the final subject-mode matrix:

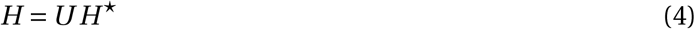

The mMIGP approach is equivalent to performing an approximate PCA on feature-concatenated data. The advantage is that it does not need to fit all data into the memory, and even can be parallelized across modalities^18^. This approach is also equivalent to applying mCCA across all modalities^40^.

### Sparse dictionary learning for voxel-space dimension reduction

If the resolution of the data is high and the number of modalities is large, applying just the mMIGP reduction still leaves FLICA as being memory and computationally expensive. Therefore, we propose a method that can effectively reduce the voxel dimension, and preserve the important spatial information for subsequent FLICA spatial modelling. Although the most obvious ways of voxel subsampling are either to apply regular spatial down-sampling (similarly, local voxel clustering) or apply PCA *within each modality*, the former only focuses on the local patterns^22^ (and does not adapt downsampling to local variations in redundant information across voxels) while the later empirically finds more global and noise patterns in neuroimaging data, and does not work at all well empirically in this context (see also Allen, et al.^23^ and references therein).

The method we used here is sparse Dictionary Learning (DicL)^19^, which effectively performs ‘voxel grouping’ in both local and global fashion. It can be used directly on each of the original z-score normalized modalities, i.e., *Y*^(*k*)^,*k* = 1,2,…,*K*, or on the mMIGP reduced data, i.e., *V*^(*k*)^,*k* = 1,2,…,*K*. Taking the former as an example, the sparse DicL is adopted here:

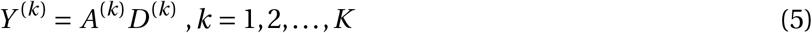

where *D*^(*k*)^ is *sparse spatial dictionary basis*, and *A*^(*k*)^ is the *feature loadings* with each column representing a linear combination of information from a group of voxels which might either be a local cluster or spatially distributed network. By minimizing an *l*_1_-regularized sparse-coding objective function, a local optimal solution can be obtained:

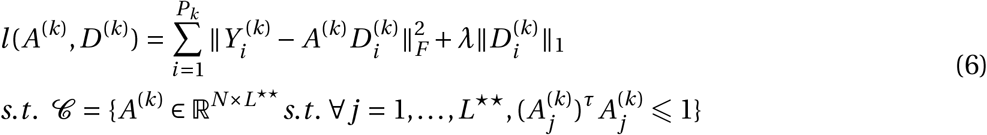

where subscript *i* represents the *i*-th column of the corresponding matrix, and *λ* is a regularization parameter. The *l*_1_-regularization term enforces that the learned spatial loadings *D*^(*k*)^ are sparse. The objective function can be efficiently optimized by a block-coordinate descent optimizer with warm restarts. It has been implemented in the SPAMS package (http://spams-devel.gforge.inria.fr/). Compared with simply using PCA in this step, sparse DicL has three advantages: (1) the spatial loading matrix *D*^(*k*)^ can be sparse, so a smaller number of voxels are involved in each column of the dictionary; (2) the columns of the dictionary do not need to be orthogonal to each other, which is more flexible; (3) an “overcomplete” dictionary is allowed, i.e., the number of dictionary basis vectors can exceed the minimum of *N* and *P_k_*, which further increases the flexibility.

After the above modality-wise DicL, the final inputs to FLICA are the matrices *A*^(*k*)^, *k* = 1,2,…, *K*, of size *N* × *L*^★★^ if we use *Y*^(*k*)^, *k* = 1,2,…, *K*, or *L*^★^ × *L*^★★^ if we use *V*^(*k*)^, *k =* 1,2,…, *K*. Note that (unlike the typical approach of feeding spatial PCA eigenvectors into ICA) we are not feeding the spatial dictionary basis (*D*^(*k*)^) into the FLICA core modelling, but the feature loadings (*A*^(*k*)^). To get the spatial loading matrices from FLICA, we do voxel-wise multiple regression where the target variable is a voxel and the design matrix is the FLICA subject mode. We could change the order by applying DicL first and then mMIGP, but this empirically has a lower computation efficiency.

### Evaluation of BigFLICA in simulations

We simulated 500 subjects, and each had two modalities, which were both 30 × 30 × 30 images. We first simulated *K* ground-truth (independent) spatial maps *X*; each of these was a 30 × 30 × 30 image. The spatial maps were a weighted sum of two Gaussian white noise images, where the first one was 30 × 30 × 30 with weight 0.05, and the second was a 5 × 5 × 5 cube randomly located in the full image with weight 0.95. Then, random positive component weights *W*, Gaussian random subject loadings *H* and Gaussian white noise terms *E* were simulated. Finally, after vectorizing each spatial map and noise term, the data for a single modality *Y* was generated as *Y* = *HWX* + *σE*, where *σ* was a parameter to control the signal-to-noise ratio (SNR). A small amount of spatial smoothing using a Gaussian kernel was applied to spatial maps *X* and noise terms *E* to mimic real image data. Each of the two modalities also had 5 unique spatial maps that were not shared by each other. The voxels were z-score normalized before feeding into the subsequent FLICA analysis. The SNR was defined as: *var*(*vec*(*HW X*))/*var*(*vec*(*σE*)).

Performance evaluation: When FLICA was applied to the simulated data, the number of independent components was always set to the ground truth *K*. The performance was measured by the similarity between estimated subject-mode matrix *H*^★^ and the ground truth *H*. The similarity was measured by the greedy matching of the components based on maximum correlation and then estimating the mean correlations across components.

Evaluation of mMIGP for subject-space dimension reduction: After generating simulated data, we reduced the data to varying dimensions (*L*^★^ = 50, 100, 200, 300, 400) using mMIGP, and then fed the reduced data into FLICA. This was compared with the original FLICA. The number of ground-truth components was set to 25,35,45 and the SNR was set to 4, 1, 0.25, 0.04. All simulations were repeated 50 times.

Evaluation of DicL for voxel-space dimension reduction: To evaluate the influence of the DicL parameters on the subsequent FLICA results, we performed the DicL on simulated data using varying parameter combinations (*Λ* = 0.1 to 16 and *L*^★★^ = 100 to 3000) followed by FLICA (nIC = 25, 50, 100). This was compared with the original FLICA. The SNR was set to 4,1,0.25,0.04, and the number of iterations for the DicL was set to 50, because we empirically find that this number of iterations is sufficient for DicL to converge to a stable result in simulation and real data. All simulations were repeated 50 times.

### HCP and UK Biobank data

The voxel/vertex-wise neuroimaging data of 81 different modalities of 1,003 subjects from the HCP S1200 data release were used in this paper^2^. The preprocessing was conducted by the HCP team using an optimized pipeline^41^. The 81 modalities included (1) 25 resting-state ICA dual-regression spatial maps (z-score normalized); (2) 47 unique task contrast maps as z-statistics from 7 different fMRI tasks; (3) 3 T1-image derived modalities (grey matter volume, surface area, surface thickness);(4) 6 Tract-Based Spatial Statistics (TBSS) features from diffusion MRI (FA, L1, L2, L3, MD, MO)^42^. In addition, 158 nIDPs were used here, which was the same as our previous study^43^. Names of nIDPs are in **Supplementary File 1**.

The UK Biobank imaging data were mainly preprocessed by FSL^34,35^ and FreeSurfer^44^ following an optimized pipeline^45^ (https://www.fmrib.ox.ac.uk/ukbiobank/). The voxel-wise neuroimaging data of 47 modalities of 14,053 subjects were used in this paper, including: (1) 25 “modalities” from the resting-state fMRI ICA dual-regression spatial maps (z-score normalized); (2) 6 modalities from the emotion task fMRI: 3 contrasts (shapes, faces, faces>shapes) of z-statistics and 3 *contrasts of parameter estimate* maps; (3) 10 diffusion MRI derived modalities (9 TBSS features, including FA, MD, MO, L1, L2, L3, OD, ICVF, ISOVF^42,46^ and a summed tractography map of 27 tracts from AutoPtx in FSL); (4) 4 T1-MRI derived modalities (grey matter volume and Jacobian map (which shows expansion/contraction generated by the nonlinear warp to standard space, and hence reflects local volume) in the volumetric space, and cortical area and thickness in the Freesurfer’s fsaverage surface space); (5) 1 susceptibility-weighted MRI map (T2-star); (6) 1 T2-FLAIR MRI derived modality (white matter hyperintensity map estimated by BIANCA47). A detailed description is in **Table. S5**. In addition, the 8,787 nIDPs were included, butwe retained the 7,245 of those, that have at least 1,000 non-missing values (subjects). Names of nIDPs are in **Supplementary Files**. Group-level resting-state independent component spatial maps and task activation z-statistic maps are in the **Supplementary Files**.

When carrying out nIDP prediction, a total of 13 and 54 confounding variables were regressed out from nIDPs using linear regression in the HCP and the UKB datasets respectively (**Supplementary Materials**). Subjects with a missing modality were imputed by the mean value of all other subjects. We did not impute the missing nIDPs.

### Comparing BigFLICA with the original FLICA on real data

On real data, we do not know the ground truth components, and the data may not follow the assumptions of ICA. Therefore, we rely on the performance of predicting nIDPs as a surrogate criterion to evaluate different methods. We applied the proposed mMIGP approach to HCP data and a subset of 1,036 UKB subjects (so that the original FLICA is computationally tractable). Elastic-net regression, from the glmnet package^48^, was used to predict the nIDPs using FLICA’s subject modes as model regressors (features). This approach is widely-used and has been shown to achieve a robust and state-of-the-art performance in many neuroimaging studies^24,25^. To evaluate the model performance, for each nIDP, we used 5-fold cross validation, and compute Pearson correlation between the predicted and true values of each nIDP across the 5 test sets. As there are tuning parameters within the Elastic-net regression, in each training set, we performed a nested 5-fold cross validation to tune the model parameters, and used the best model selected in the nested 5-fold cross validation to do the prediction in the test set. When comparing any two approaches, the same training-validation-testing split was used. The prediction accuracy was quantified as the Pearson correlation between predicted and the true values of each nIDP in the test sets.

To evaluate MIGP preprocessing, we reduced the dimension to varying *L*^★^ (from 100 to 500) using MIGP first and then used FLICA to extract *L* = 50 components. The original FLICA was also applied to extract 50 components. To evaluate DicL preprocessing, we used the DicL (dictionary dimension = 2000 and sparsity parameter *λ* = 1) to reduce the data dimension of each modality followed by the FLICA to extract varying numbers of components (nIC= 25, 50, 100, 200, 300). The original FLICA was also applied to extract the same numbers of components. The prediction accuracy of BigFLICA was compared with the original FLICA applied on non-reduced data.

### Statistical significance of difference of prediction accuracy between two approaches

To compare the overall prediction accuracy of two approaches (e.g. BigFLICA with mMIGP preprocessing vs. the original FLICA), we estimate the statistical significance of the difference between the prediction correlations across nIDPs. Suppose that we have a total of *p* nIDPs, we first filter out a subset of nIDPs where both methods have low prediction accuracy (*r* < 0.1 in our analysis), resulting in *p*_1_ nIDPs. If we perform a simple paired t-test, the correlation structures among nIDPs makes the samples dependent with each other, so that the p-value is not valid. Based the fact that a paired t-test is a special case of general linear model (where the *y* variable is the difference of the prediction accuracy, and the *x* variable is a column of ones, and the statistical significance is the significance of the coefficient of *x*), we used a weighted least square approach (by lscov function in Matlab) to get a reliable statistical significance estimation by taken the covariance structures between nIDPs (which is estimated as the covariance of the nIDPs-by-subject matrix) into account.

### Parameter settings of running BigFLICA in the full HCP and UKB datasets

We applied BigFLICA approach to extract varying number of target components in two datasets. In HCP, we used FLICA with DicL preprocessing only (dictionary dimension 2000 and *λ* = 1). In UKB, we used FLICA with both mMIGP and DicL preprocessing (dictionary dimension 5000, *λ* = 1 and mMIGP dimension 1000 (*>*95% explained variance)). The number of FLICA VB iterations is 1000.

### Comparing BigFLICA with multiple independent single-modality ICA decomposition

ICA is a widely-used approach for decomposing single-modality neuroimaging data, including functional MRI^43^ but also in structural MRI^49^ and diffusion MRI^50^. A natural question arises whether BigFLICA is able to combine multimodal information more effectively than the single-modality approaches such as ICA (we used the fastICA algorithm^51^), which ignores inter-modality relationships.

We first performed ICA on each modality of HCP and UKB data separately to extract 25,100, 250, 500 and 750 components. For a given component number, we built a prediction model using the concatenated ICA subject modes (across modalities) to predict each of the nIDPs. To be fair, for BigFLICA, we extract the same number of ICs to build the prediction model. For example, in the UKB data and a 25-dimensional decomposition, the predictor is a Subject×(25 × 47) matrix for single-modality ICA, where 25 is the number of components in each modality and 47 is the total number of modalities. For BigFLICA, the predictor is a Subject× 25 matrix. This is arguably a fair comparison because each of the BigFLICA modes potentially contains information from all modalities. The method to build a predictive model and evaluate this is the same as above, except that when we used the concatenated ICA subject modes, we added a univariate screening step in the training set to select the top 300 most informative features according to their correlation with an nIDP in the training set. This step, in general, boosts the predictive accuracy because the dimensionality of concatenated ICA modes is usually very high, so that many of the modes are pure noise with respect to any given nIDP. Therefore, the univariate screening can help the elastic-net regression to filter out noisy features effectively. We did not perform univariate screening when using the BigFLICA subject modes to predict nIDPs.

Besides the main results, in **Fig. S1**, we also compared, in the UKB data, the 750-dimensional BigFLICA decomposition with the 25-dimensional ICA decomposition concatenated across modalities, i.e., we have 25 × 47 features in the single-modality ICA. In this comparison, the number of features for the two methods are almost the same, but we can see that BigFLICA clearly outperforms the single-modality ICA.

### Comparing BigFLICA with hand-curated imaging-derived phenotypes

A popular choice of data analysis strategy is to extract imaging features based on expert knowledge (e.g., regional volumes and thickness, and resting-state functional connectivities between brain regions), often referred to as IDPs^1^. Brain IDPs have been shown to genetically correlate with many SNPs in our previous genome-wide association study (GWAS) in UK Biobank^5^, and they have been shown to change in many psychiatric diseases^26–28^.

We extracted 5,812 IDPs from the HCP, including (1) 199 structural MRI features from Freesurfer as provided by the HCP; (2) 4700 regional mean task activations from 47 independent task contrasts using a 100-dimensional parcellation atlas^52^; (3) 625 functional connectivities (FCs) based on a 25-dimensional ICA parcellation with partial correlation to estimate FCs; (4) 288 regional mean TBSS features (FA, L1, L2, L3, MD, MO) using the Johns Hopkins University tract atlas. The names of these IDPs are given in the **Supplementary File 3**.

We used 3,913 IDPs from UKB, including global and local features from the 6 imaging modalities (T1, T2-FLAIR, swMRI, tfMRI, rfMRI, and dMRI)^53^. The names of these IDPs are given in the **Supplementary File 4**.

We built prediction models that use IDPs or BigFLICA modes to predict each of the nIDPs using the same strategy as above. The FLICA dimension is set to 25, 100, 250, 500, 750. In addition, we also concatenated IDPs and each of the BigFLICA subject modes together to predict the nIDPs, and the performance is compared with using IDPs alone. We used a univariate screening step to select the top 300/500 most informative IDPs according to their correlation with an nIDP in the inner-fold (i.e., training set) of HCP/UKB. Finally, we also built models that use IDPs to predict each of the FLICA subject modes and vice versa, aiming to evaluate the shared variances between features extracted by these two different approaches in the same data.

### Reproducibility of BigFLICA

To test whether BigFLICA’s spatial independent components are estimated reliably, the whole UKB dataset was divided into two parts: the first part contained 7,000 subjects and the second part contained the remaining 7,503 subjects. We applied BigFLICA to the two parts separately. After estimating the subject modes, we reconstructed the z-score normalized (voxel-wise) spatial maps of each modality by regressing the subject mode against the mMIGP-reduced data. The spatial independent components of each modality were concatenated spatially and greedily paired, based on the absolute correlation between two runs. When we computed the correlations, only voxels whose absolute z-scores that are both larger than 3 in two runs were preserved (to reduce noise, given that there are huge numbers of empty voxels across all modalities for a given FLICA component in general; this does not bias the metric of reproducibility towards finding common similar patterns). **Fig. S5** (left) shows that the FLICA components have very high reproducibility in the split-half test across a varying number of components.

### Comparing BigFLICA with mCCA

We tested whether BigFLICA (independent components-based spatial modelling) was better than mCCA (eigendecomposition based modelling, which could be considered to be similar to the output of BigFLICA without running the final core FLICA unmixing - note that to enable mCCA to run requires the same mMIGP initial processing that we have added in this work) in three ways. The number of extracted components was the same when performing this comparison. First, for the prediction accuracy of nIDPs, **Fig. S3** shows that, in the UKB data, BigFLICA has a (very slightly) improved prediction accuracy compared with mCCA. Then, we proposed a hypothesis that modes of BigFLICA are more parsimonious features of nIDPs compared with mCCA, or in other word, a smaller number of modes of BigFLICA can predict the nIDPs. Results shown in **Fig. S4** validate this hypothesis: for a given number of components and a given nIDP, BigFLICA modes have a (on average) higher proportion of zero weights in the elastic-net predictions, when compared with mCCA modes. The advantage is that a more parsimonious representation usually has a better biological interpretability. Finally, we estimated and compared the split-half reproducibility of BigFLICA and mCCA. As shown in **Fig. S5** (right), BigFLICA has a much higher between-subject reproducibility than mCCA.

### Contribution of different modalities in a BigFLICA decomposition

Besides using BigFLICA for exploring the relationships between imaging and non-imaging phenotypic and behavioural data, we can also use it to investigate the relationship between different modalities. For each mode, BigFLICA estimates a vector of positive numbers reflecting the contributions of different modalities (i.e., the diagonal of each *W*_(*k*)_, where the higher the number, the more important is one modality to a mode). We concatenated all such vectors across all modes so that it is a mode-by-modality matrix *W*, and normalized each column to sum to one. Six examples of such matrices are shown in **Fig. S7**, with different numbers of estimated modes in the UKB dataset.

We then calculate each row’s sum (across columns) in *W*, thereby reflecting the overall contribution of each modality in the BigFLICA decomposition. As shown in **Fig. S8**, across all FLICAdimension-alities (numbers of estimated modes), each of the 25 resting-state fMRI dual-regression spatial maps usually has a low overall contribution, followed by task fMRI maps, while modalities reflecting more about structure of the brain (e.g., structural MRI and diffusion MRI) generally have high overall contributions. The relative differences of modality contribution between functional MRI-related modalities and structural/diffusion MRI-related modalities become larger with increasing number of estimated modes. We further estimated the total shared variances between a lower dimensional BigFLICA decomposition and a higher dimensional decomposition. **Table S4** shows that a higher dimensional decomposition explains almost all variances of a lower dimensional decomposition (upper triangle of the table), while a lower dimensional decomposition can explain a large proportion of the variances of a higher dimensional decomposition.

### Relationship between different modalities in a BigFLICA decomposition

We calculated the cosine similarity between different columns of *W* (using the 750-dimensional BigFLICA decomposition), to measure the similarity of different modalities in terms of their contribution to the BigFLICA decomposition, i.e., the more similar information two modalities carry, the more likely they will have similar contribution to a mode. **Fig. S9a** shows that the modality relationship matrix is clearly grouped into three large clusters. The first is all resting-state modalities, while the second is the task fMRI maps, and the third is the diffusion MRI, structural MRI-related modalities and swMRI. The white matter hyperintensity map (T2 lesions) forms a single cluster. As a comparison, we also performed a 50-dimensional ICA decomposition within each modality, and calculated the shared variances between every pair of 50 ICs in two modalities using a simple multivariate regression model. As shown in Fig. S9b, we also observed a similar pattern as Fig. S9a. The main difference is that in Fig. S9a, there are relatively stronger correlations within resting-state modalities and between resting-state and other modalities, but weaker correlations between task modalities and structural related modalities. These results reflect the fact that the multimodal modelling effects of BigFLICA learn different inter-modality relationships compared with single-modality ICA.

## Supplementary Materials

### Confounding variables regressed out in our analysis

UKB dataset: age, age squared, age X sex, age squared X sex, age (quantile normalised), age squared (quantile normalised), age X sex (quantile normalised), age squared X sex (quantile normalised), rfMRI head motion, tfMRI head motion, head size scaling, rfMRI head motion squared, tfMRI head motion squared, [4] confounds relating to bed position in scanner (x), [4] confounds relating to bed position in scanner (y), [4] confounds relating to bed position in scanner (z), [4]confounds relating to bed position in scanner (table), [4]confounds relating to bed position in scanner (x) squared, [4] confounds relating to bed position in scanner (y) squared, [4] confounds relating to bed position in scanner (z) squared, [4]confounds relating to bed position in scanner (table) squared, [10] confounds modelling slow date-related drift 1, [10] confounds modelling slow date-related drift 2, [10] confounds modelling slow date-related drift 3, [10] confounds modelling slow date-related drift 4, [10] confounds modelling slow date-related drift 5, [10] confounds modelling slow date-related drift 6, [10] confounds modelling slow date-related drift 7, [10] confounds modelling slow date-related drift 8, [10] confounds modelling slow date-related drift 9, [10] confounds modelling slow date-related drift 10, rfMRI head motion (quantile normalised), tfMRI head motion (quantile normalised), head size scaling (quantile normalised), [4] confounds relating to bed position in scanner (x) (quantile normalised), [4] confounds relating to bed position in scanner (y) (quantile normalised), [4] confounds relating to bed position in scanner (z) (quantile normalised), [4] confounds relating to bed position in scanner (table) (quantile normalised), [4] confounds relating to bed position in scanner (x) squared (quantile normalised), [4] confounds relating to bed position in scanner (y) squared (quantile normalised), [4] confounds relating to bed position in scanner (z) squared (quantile normalised), [4] confounds relating to bed position in scanner (table) squared (quantile normalised), [10] confounds modelling slow date-related drift 1 (quantile normalised), [10] confounds modelling slow date-related drift 2 (quantile normalised), [10]confounds modelling slow date-related drift 3 (quantile normalised), [10]confounds modelling slow date-related drift 4 (quantile normalised),[10]confounds modelling slow date-related drift 5 (quantile normalised), [10] confounds modelling slow date-related drift 6 (quantile normalised), [10] confounds modelling slow date-related drift 7 (quantile normalised), [10] confounds modelling slow date-related drift 8 (quantile normalised), [10] confounds modelling slow date-related drift 9 (quantile normalised), [10]confounds modelling slow date-related drift 10 (quantile normalised), imaging centre, sex.

HCP dataset: image reconstruction version, age, age squared, sex, age X sex, age squared X sex, race, ethnicity, rfMRI motion, Height, Weight, FS_IntraCranial_Vol, FS_BrainSeg_Vol.

**Figure S1:**
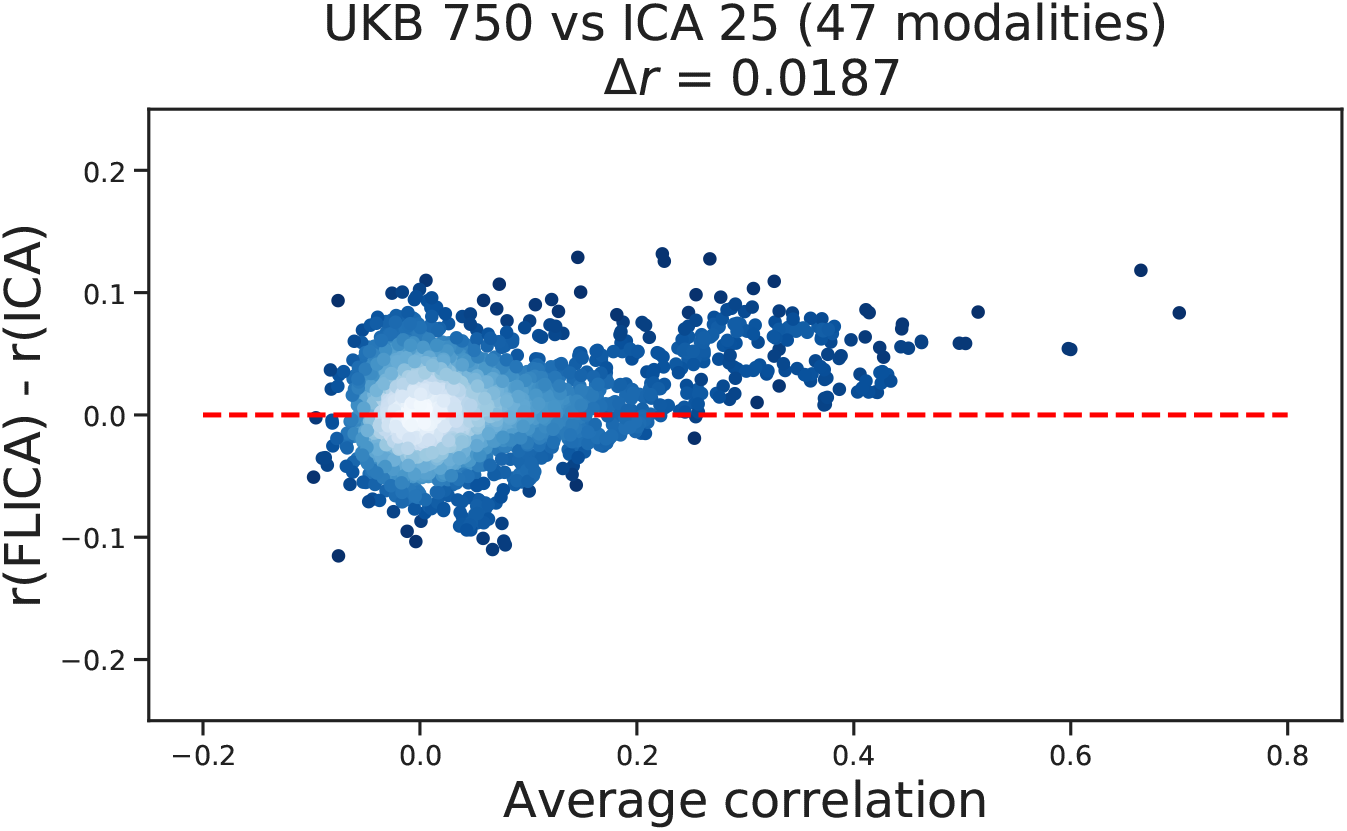
Comparing the prediction performance of the 750-dimensional FLICA with the 25-dimensional single-modality ICA concatenated across 47 modalities in the UKB data.

**Figure S2:**
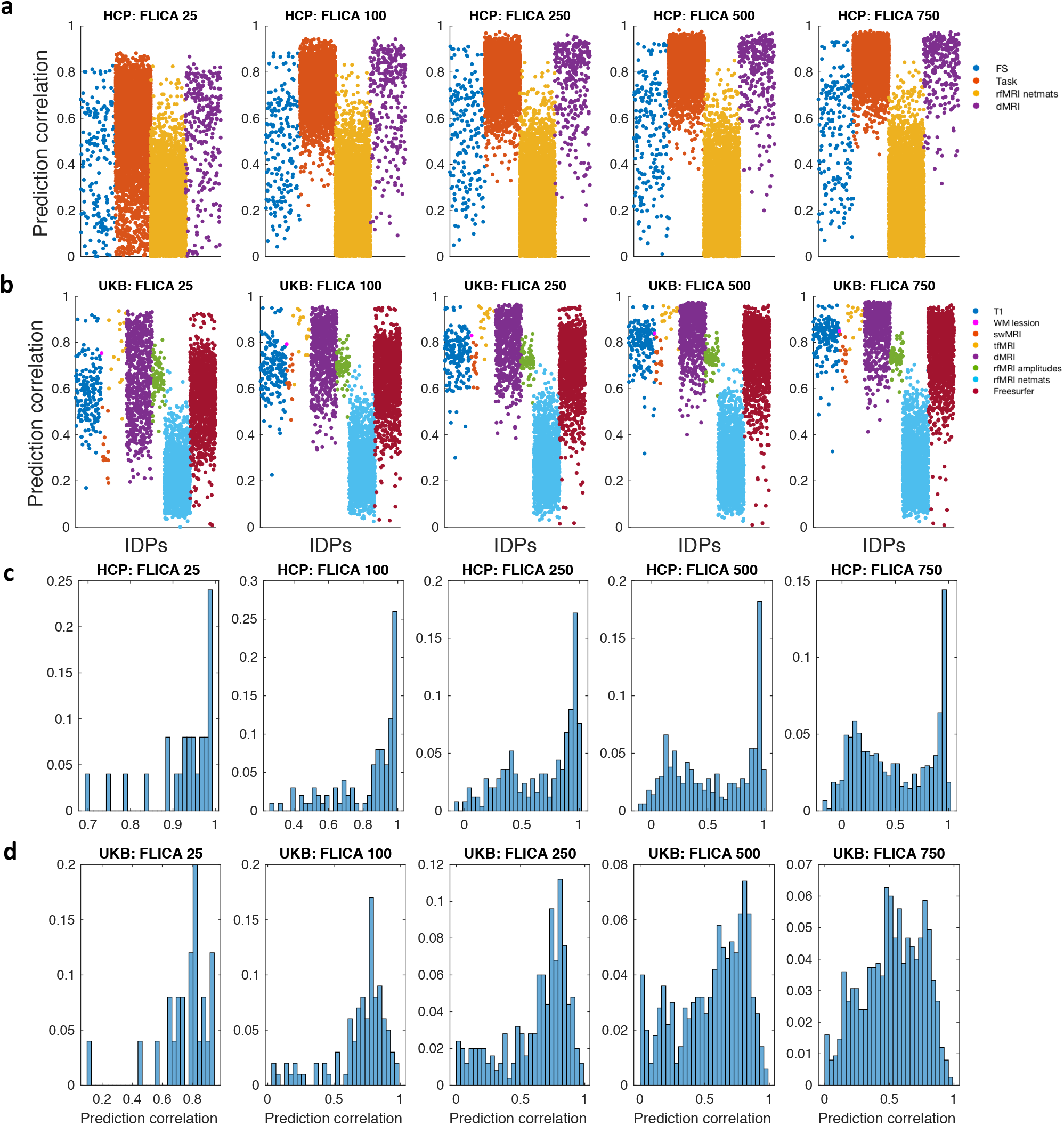
Relationships between FLICA and IDPs. (**a,b**) The plots show the results of predicting each IDP using BigFLICA modes in **a** the HCP and **b** the UKB dataset. The IDPs are appearing in order along the x axis, and are grouped and coloured by modality types. (**a,b**) The histograms of predicting BigFLICA modes using all IDPs in **c** the HCP and **d** the UKB dataset.

**Figure S3:**
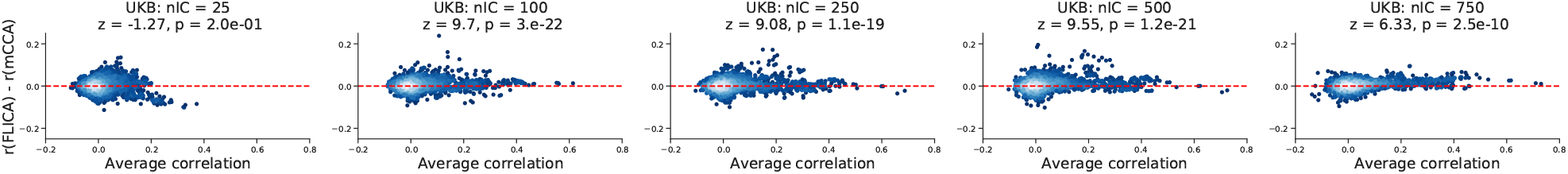
Comparing FLICA and mCCA in the UKB data. Comparing the predictive performance of FLICA with mCCA (or equivalently the subject-by-component matrix obtained in the mMIGP step) across different numbers of extracted components in the UKB dataset. The FLICA and mCCA dimensions are the same in each figure.

**Figure S4:**
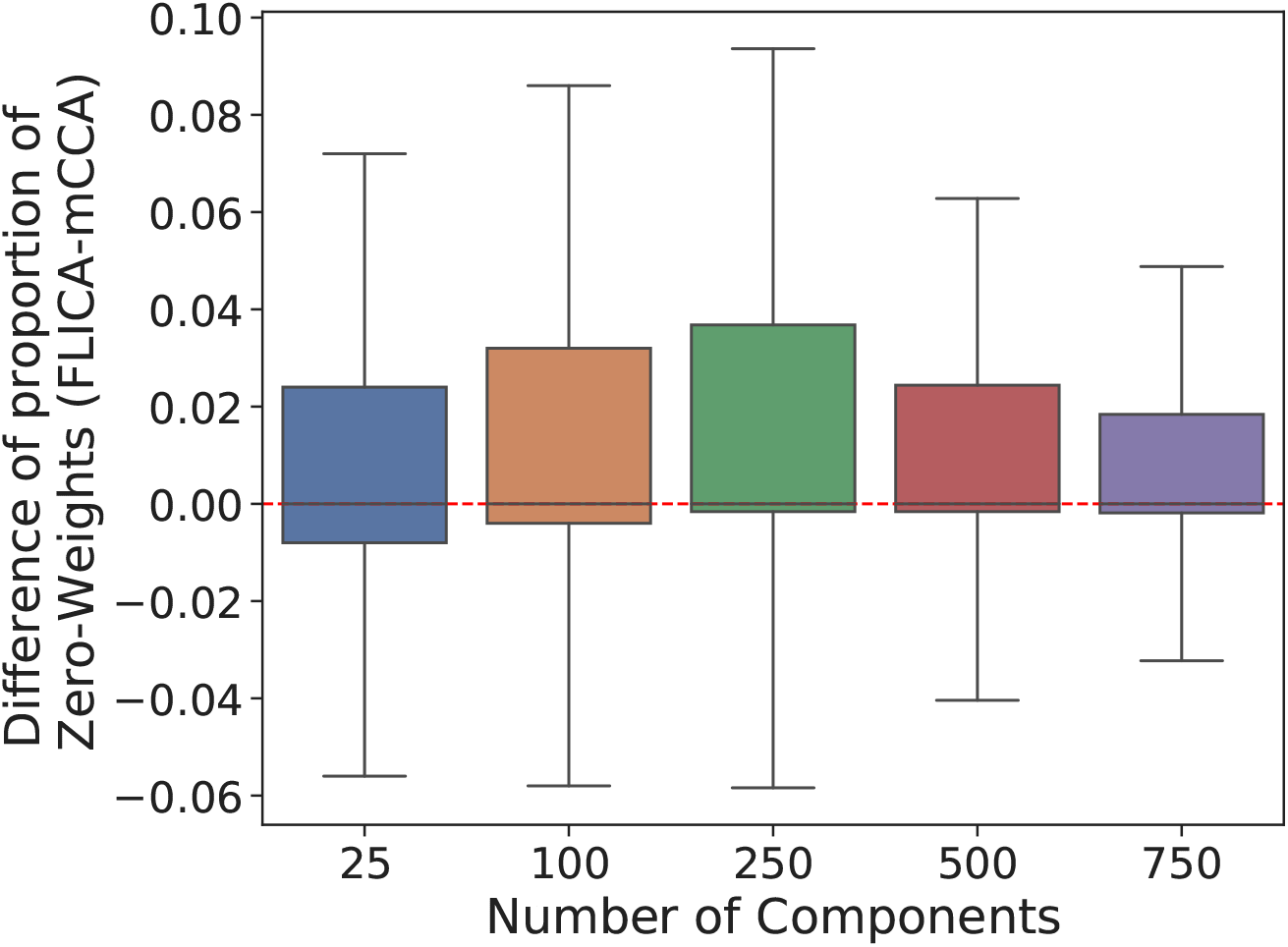
The difference of the proportion of zeros weights (BigFLICA-mCCA) in predicting nIDPs across 5 dimensions of decomposition in the UKB data.

**Figure S5:**
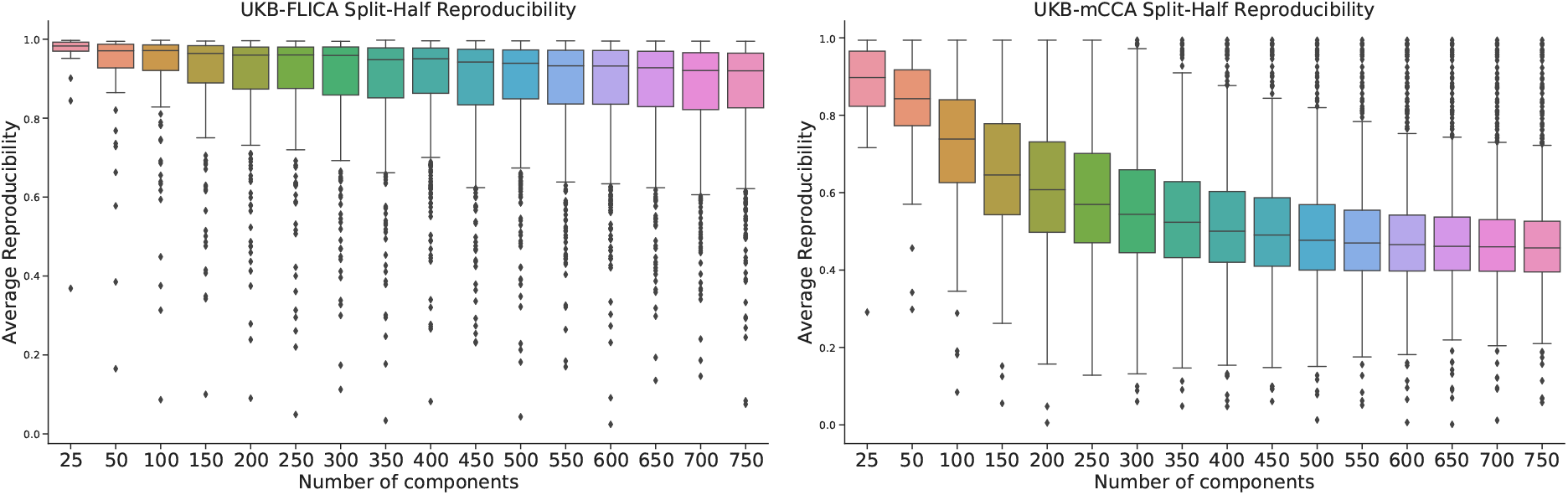
Split-half reproducibility of FLICA and mCCA spatial maps in the UKB dataset. The split-half reproducibility of BigFLICA and mCCA in the UKB dataset by first computing the correlation between modality-wise concatenated spatial maps after eliminating low-weight voxels and then greedy matching.

**Figure S6:**
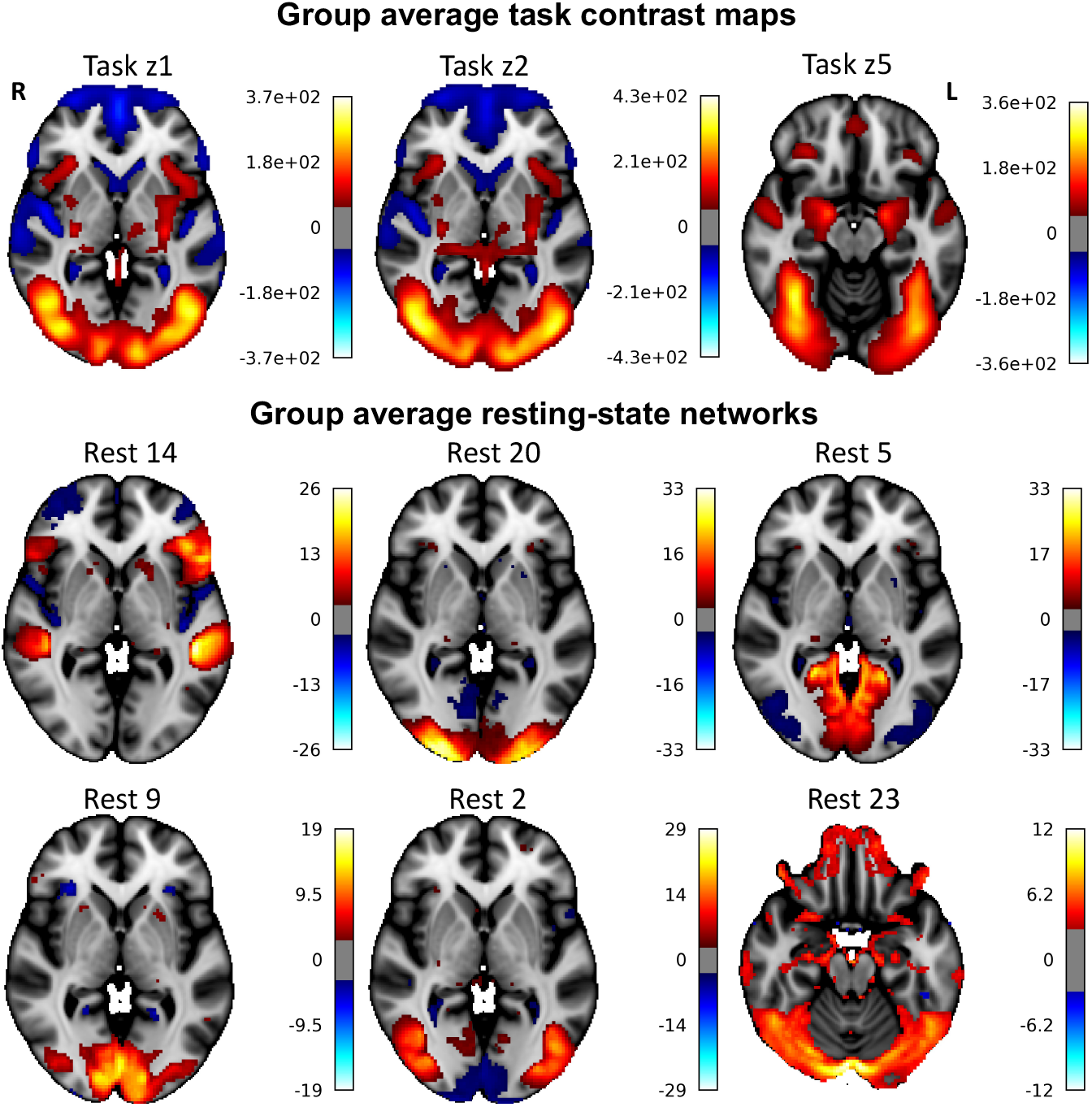
Group average maps of task activations and resting-state networks in the UKB dataset. These are provided to help interpret the population variability maps (modulations of these maps) shown in **Fig. 5**. Top: group average of emotion task activation z-statistic maps (task z1: “shapes”, task z2: “face”, task z5: “faces>shapes”). Group average task contrast (effect size) maps c1, c2 and c5 are highly similar to z-stat maps so that they are not shown. Bottom: group average resting-state networks from a 25-dimensional ICA parcellation in the UKB data. The six maps shown here are the networks from **Fig. 5**.

**Figure S7:**
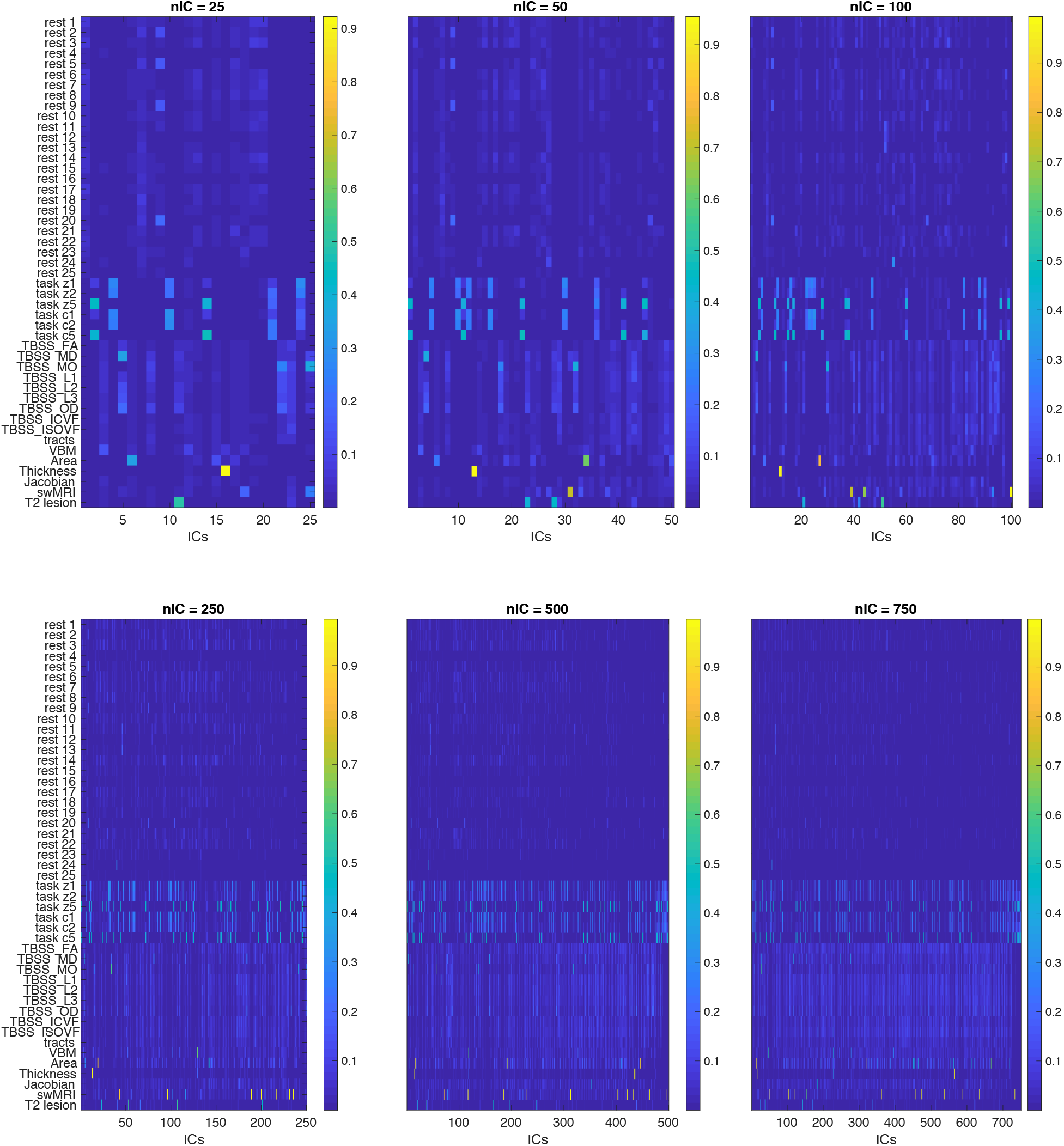
The contribution of each modality in each BigFLICA mode (independent component).

**Figure S8:**
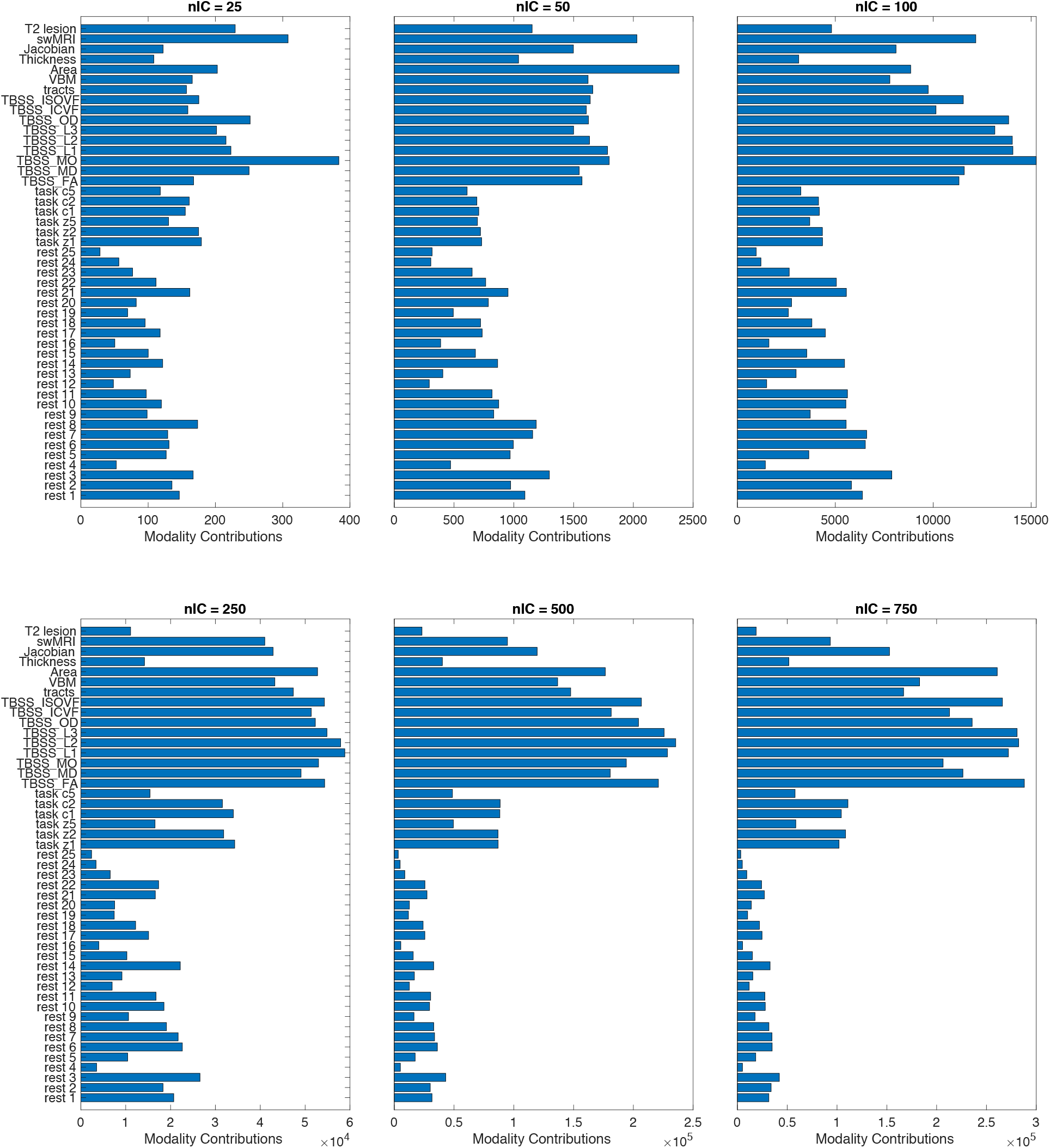
The relative contribution of different modalities of a BigFLICA decomposition (nIC=25-750) in the UKB data. For each modality, we take the sum of its overall contribution (estimated by BigFLICA) across all components.

**Figure S9:**
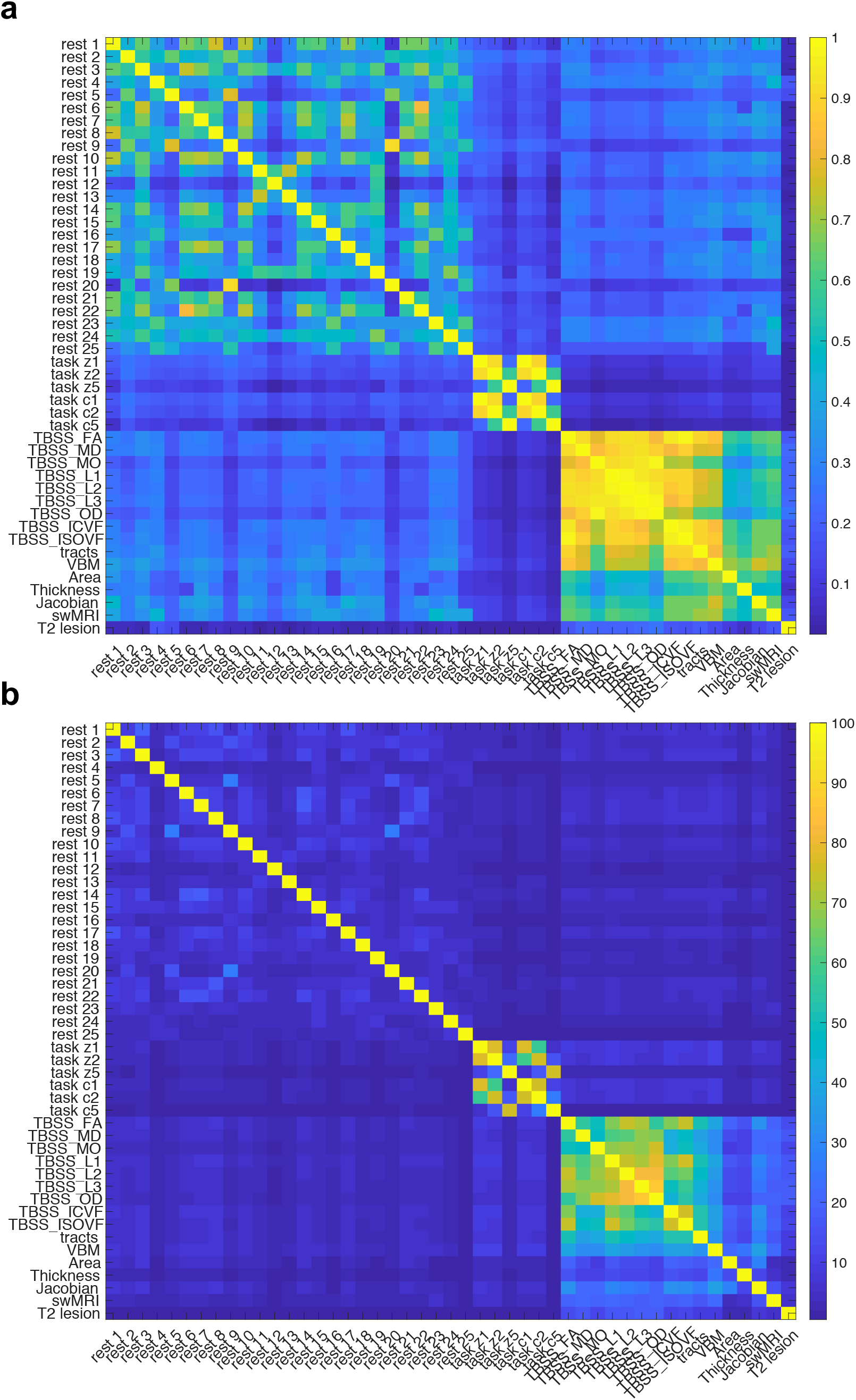
The relationships between different modalities in the UKB data. a). The cosine similarity of modality contributions across 750 components (estimated by BigFLICA) between every pair of modalities. b). The amount of shared variance between two 50-dimensional single-modality ICA decompositions in each pair of modalities.

**Table S1:** Comparison of prediction performance of **Cognitive Phenotypes** and **Health and Medical History Health Outcomes** between BigFLICA (nIC=750) and 3,913 IDPs in the UKB dataset. We excluded an nIDP if both methods have prediction r-value< 0.1.

| Variable Names | r-value FLICA | -log10(p) FLICA | r-value IDPs | -log10(p) IDPs | Percent Improvement | Sample size |
| --- | --- | --- | --- | --- | --- | --- |
| **cognitive phenotypes** |  |  |  |  |  |  |
| Digits entered correctly (0.1) | 0.104 | 3.6 | 0.003 | 0.3 | 3141.6 | 1126 |
| Time to complete round (1.2) | 0.108 | 11.7 | 0.073 | 5.9 | 47.4 | 4119 |
| Number of fluid intelligence questions attempted within time limit (1.0) | 0.107 | 11.6 | 0.078 | 6.6 | 37.6 | 4116 |
| Duration to first press of snap-button in each round (2.7) | 0.11 | 37.6 | 0.081 | 21.1 | 35.1 | 13749 |
| Mean time to correctly identify matches (0.0) | 0.108 | 38.6 | 0.083 | 23.1 | 30.6 | 14471 |
| Duration screen displayed (2.0) | 0.103 | 33.6 | 0.079 | 20.3 | 30.2 | 13809 |
| Duration to first press of snap-button in each round (2.10) | 0.104 | 33.7 | 0.08 | 20.3 | 30.1 | 13751 |
| Duration to complete alphanumeric path (trail #2) (0.0) | 0.145 | 33.7 | 0.113 | 20.8 | 28.7 | 6966 |
| Time to complete round (0.2) | 0.113 | 41 | 0.088 | 25.5 | 27.8 | 14115 |
| Number of symbol digit matches made correctly (0.0) | 0.156 | 43.5 | 0.122 | 27 | 27.7 | 7862 |
| Number of symbol digit matches attempted (0.0) | 0.167 | 49.8 | 0.132 | 31.4 | 26.6 | 7862 |
| Number of fluid intelligence questions attempted within time limit (2.0) | 0.151 | 68.4 | 0.122 | 44.9 | 23.8 | 13362 |
| Duration to first press of snap-button in each round (2.11) | 0.1 | 31.6 | 0.082 | 21.3 | 22.9 | 13741 |
| Time to complete round (0.1) | 0.143 | 36 | 0.117 | 24.4 | 22.5 | 7670 |
| Time to complete round (2.2) | 0.114 | 39.4 | 0.093 | 26.8 | 22.1 | 13394 |
| Mean time to correctly identify matches (2.0) | 0.141 | 61.6 | 0.116 | 42 | 21.6 | 13768 |
| Maximum digits remembered correctly (0.0) | 0.15 | 38.2 | 0.129 | 28.7 | 15.9 | 7465 |
| Fluid intelligence score (2.0) | 0.256 | 198.7 | 0.232 | 162.6 | 10.3 | 13362 |
| Fluid intelligence score (0.0) | 0.199 | 47.8 | 0.183 | 40.5 | 8.8 | 5266 |
| Touchscreen duration (2.0) | 0.127 | 52.5 | 0.121 | 48 | 4.7 | 14412 |
| Fluid intelligence score (1.0) | 0.18 | 31 | 0.188 | 33.5 | -3.9 | 4116 |
| Number of fluid intelligence questions attempted within time limit (0.0) | 0.098 | 12.4 | 0.103 | 13.4 | -4 | 5266 |
| Fluid intelligence score (0.0) | 0.198 | 72.1 | 0.209 | 80.5 | -5.3 | 8090 |
| Touchscreen duration (1.0) | 0.103 | 10.8 | 0.127 | 16 | -19.2 | 4134 |
| **Health and Medical History Health Outcomes** |  |  |  |  |  |  |
| Age asthma diagnosed (0.0) | 0.122 | 5.6 | 0.051 | 1.5 | 139.4 | 1397 |
| Interpolated Age of participant when non-cancer illness first diagnosed (0.3) | 0.102 | 5.1 | 0.072 | 2.9 | 42.7 | 1794 |
| Interpolated Year when non-cancer illness first diagnosed (0.3) | 0.104 | 5.3 | 0.074 | 3 | 41.5 | 1794 |
| Medication for cholesterol, blood pressure, diabetes, or take exogenous hormones (2.0) | 0.153 | 40.4 | 0.113 | 22.2 | 36.4 | 7507 |
| Treatment/medication code (1140884600 - metformin) | 0.121 | 48.1 | 0.092 | 28.1 | 32 | 14503 |
| Number of treatments/medications taken (0.0) | 0.13 | 55.6 | 0.099 | 32.4 | 31.8 | 14503 |
| Age started wearing glasses or contact lenses (2.0) | 0.158 | 73.8 | 0.124 | 45.8 | 27.3 | 13129 |
| Number of self-reported non-cancer illnesses (0.0) | 0.115 | 43.7 | 0.093 | 28.6 | 24.4 | 14503 |
| Number of treatments/medications taken (1.0) | 0.13 | 16.7 | 0.107 | 11.7 | 21.4 | 4134 |
| Age started wearing glasses or contact lenses (0.0) | 0.149 | 61.6 | 0.124 | 43.2 | 19.8 | 12321 |
| Number of treatments/medications taken (2.0) | 0.188 | 114.4 | 0.157 | 80.2 | 19.4 | 14435 |
| Treatment/medication code (1140879802 - amlodipine) | 0.113 | 41.9 | 0.096 | 30.5 | 17.8 | 14503 |
| Medication for cholesterol, blood pressure or diabetes (0.0) | 0.15 | 35.1 | 0.131 | 26.7 | 15.1 | 6756 |
| Diabetes diagnosed by doctor (2.0) | 0.124 | 49.6 | 0.108 | 38.3 | 14.2 | 14379 |
| Non-cancer illness code, self-reported (1220 - diabetes) | 0.109 | 39.1 | 0.098 | 31.8 | 11.1 | 14503 |
| Diagnoses - secondary ICD10 (I10 - I10 Essential (primary) hypertension) | 0.146 | 69.8 | 0.135 | 59.5 | 8.4 | 14503 |
| Overall health rating (0.0) | 0.113 | 41.6 | 0.104 | 35.7 | 8.1 | 14477 |
| Non-cancer illness code, self-reported (1065 - hypertension) | 0.218 | 155.7 | 0.203 | 133.6 | 7.9 | 14503 |
| Medication for cholesterol, blood pressure or diabetes (2.0) | 0.145 | 33.2 | 0.136 | 29.3 | 6.7 | 6829 |
| Overall health rating (2.0) | 0.11 | 39.5 | 0.113 | 41.7 | -2.8 | 14388 |
| Non-cancer illness code, self-reported (1261 - multiple sclerosis) | 0.111 | 40.2 | 0.118 | 46 | -6.6 | 14503 |

**Table S2:**
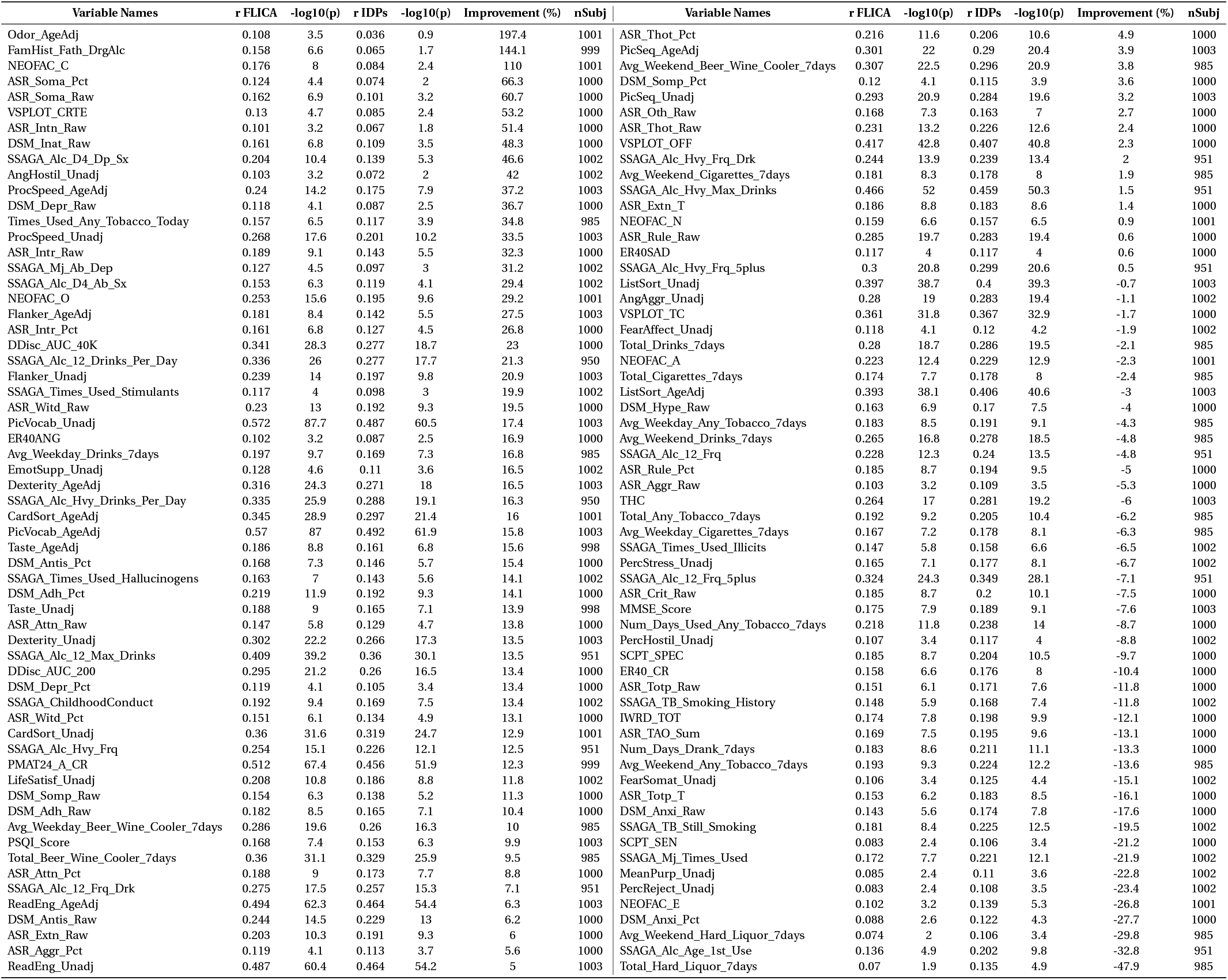
Comparison of prediction performance of 158 nIDPs between FLICA (nIC=100) and 5,812 IDPs in the HCP dataset. We excluded an nIDP if both methods have prediction r-value< 0.1. The meanings of each variables can be found at HCP wiki: https://wiki.humanconnectome.org/display/PublicData/HCP+Data+Dictionary+Public-+Updated+for+the+1200+Subject+Release

**Table S3:** Three examples of top 10 most significant correlations of BigFLICA modes (left) and IDPs (right) with nIDPs in UKB dataset.

| BigFLICA modes | r-value | p-value | IDP names | r-value | p-value |
| --- | --- | --- | --- | --- | --- |
| **Top 10 modes/IDPs correlate with *fluid intelligence*** |  |  |  |  |  |
| IC25 | -0.146 | 6.54E-65 | IDP_tfMRI_90th-percentile_BOLD_shapes | -0.074 | 1.05E-15 |
| IC57 | -0.122 | 3.49E-45 | IDP_tfMRI_median_BOLD_shapes | -0.069 | 1.28E-13 |
| IC332 | -0.081 | 7.26E-21 | IDP_tfMRI_90th-percentile_zstat_faces-shapes_amygdala | 0.068 | 1.48E-13 |
| IC484 | -0.072 | 9.87E-17 | rfMRI amplitudes (ICA25 node 6) | 0.066 | 2.75E-13 |
| IC4 | -0.069 | 1.18E-15 | rfMRI connectivity (ICA100: IC13-IC32) | -0.065 | 3.82E-13 |
| IC27 | 0.064 | 1.98E-13 | IDP_tfMRI_90th-percentile_zstat_shapes | -0.063 | 7.95E-12 |
| IC188 | -0.058 | 2.80E-11 | IDP_tfMRI_median_zstat_faces-shapes | 0.063 | 1.20E-11 |
| IC708 | -0.055 | 1.51E-10 | IDP_tfMRI_median_zstat_faces-shapes_amygdala | 0.062 | 1.51E-11 |
| IC164 | 0.055 | 1.68E-10 | rfMRI connectivity (ICA100: IC11-IC19) | 0.056 | 3.94E-10 |
| IC47 | -0.054 | 3.40E-10 | IDP_tfMRI_median_BOLD_faces-shapes | 0.057 | 5.16E-10 |
| **Top 10 modes/IDPs correlate with *Age started wearing glasses or contact lenses*** |  |  |  |  |  |
| IC164 | 0.101 | 2.74E-31 | IDP_tfMRI_90th-percentile_BOLD_faces-shapes | 0.081 | 5.38E-18 |
| IC25 | 0.067 | 1.63E-14 | IDP_tfMRI_median_BOLD_faces-shapes | 0.064 | 7.16E-12 |
| IC249 | 0.057 | 5.65E-11 | IDP_tfMRI_median_zstat_faces-shapes | 0.064 | 1.01E-11 |
| IC13 | 0.054 | 7.68E-10 | rfMRI connectivity (ICA100: IC4-IC40) | -0.061 | 1.81E-11 |
| IC138 | 0.041 | 2.33E-06 | rfMRI connectivity (ICA100: IC17-IC42) | -0.060 | 4.34E-11 |
| IC190 | 0.038 | 1.65E-05 | IDP_tfMRI_90th-percentile_zstat_faces-shapes | 0.061 | 9.88E-11 |
| IC563 | -0.037 | 1.84E-05 | rfMRI amplitudes (ICA25 node 19) | 0.056 | 8.97E-10 |
| IC656 | 0.037 | 1.98E-05 | rfMRI amplitudes (ICA100 node 16) | 0.054 | 3.08E-09 |
| IC297 | -0.037 | 2.32E-05 | rfMRI connectivity (ICA100: IC15-IC43) | -0.050 | 3.14E-08 |
| IC580 | 0.037 | 2.53E-05 | rfMRI connectivity (ICA100: IC21-IC28) | 0.049 | 6.90E-08 |
| **Top 10 modes/IDPs correlate with *hypertension*** |  |  |  |  |  |
| IC259 | 0.121 | 2.95E-48 | IDP_dMRI_TBSS_MD_External_capsule_L | 0.135 | 4.17E-53 |
| IC38 | -0.111 | 3.09E-41 | IDP_dMRI_TBSS_MD_External_capsule_R | 0.132 | 6.44E-51 |
| IC319 | 0.094 | 5.98E-30 | IDP_dMRI_TBSS_L1_External_capsule_L | 0.130 | 2.29E-49 |
| IC29 | 0.091 | 5.82E-28 | IDP_dMRI_TBSS_L3_External_capsule_R | 0.126 | 6.51E-47 |
| IC40 | 0.088 | 2.82E-26 | IDP_dMRI_TBSS_L3_External_capsule_L | 0.126 | 1.70E-46 |
| IC1 | -0.083 | 1.57E-23 | IDP_dMRI_TBSS_L3_Anterior_limb_of_internal_capsule_L | 0.122 | 4.64E-44 |
| IC171 | -0.073 | 1.00E-18 | IDP_dMRI_TBSS_L1_External_capsule_R | 0.122 | 5.37E-44 |
| IC26 | -0.067 | 6.73E-16 | IDP_dMRI_TBSS_ISOVF_External_capsule_L | 0.122 | 7.49E-44 |
| IC176 | -0.062 | 6.75E-14 | IDP_dMRI_TBSS_L2_Anterior_limb_of_internal_capsule_L | 0.120 | 2.60E-42 |
| IC84 | 0.057 | 9.14E-12 | IDP_dMRI_TBSS_MD_Anterior_limb_of_internal_capsule_L | 0.118 | 2.91E-41 |
| **Top 10 modes/IDPs correlate with *handedness*** |  |  |  |  |  |
| IC235 | -0.226 | 5.71E-168 | rfMRI connectivity (ICA100: IC29-IC34) | 0.115 | 1.30E-40 |
| IC408 | -0.079 | 1.76E-21 | rfMRI connectivity (ICA25: IC1-IC6) | 0.095 | 6.93E-28 |
| IC569 | 0.066 | 1.90E-15 | rfMRI connectivity (ICA100: IC10-IC34) | 0.095 | 7.70E-28 |
| IC382 | 0.051 | 8.40E-10 | rfMRI connectivity (ICA25: IC14-IC22) | 0.085 | 7.43E-23 |
| IC251 | 0.047 | 1.13E-08 | rfMRI connectivity (ICA100: IC3-IC19) | 0.085 | 8.01E-23 |
| IC232 | 0.043 | 2.12E-07 | rfMRI connectivity (ICA100: IC14-IC34) | 0.081 | 7.28E-21 |
| IC742 | 0.042 | 4.41E-07 | rfMRI connectivity (ICA25: IC1-IC22) | 0.080 | 2.93E-20 |
| IC643 | -0.039 | 2.39E-06 | rfMRI connectivity (ICA100: IC30-IC34) | -0.074 | 1.46E-17 |
| IC419 | 0.036 | 1.66E-05 | rfMRI connectivity (ICA100: IC27-IC52) | 0.073 | 3.07E-17 |
| IC332 | 0.036 | 1.76E-05 | rfMRI connectivity (ICA100: IC6-IC13) | 0.071 | 3.39E-16 |

**Table S4:** Percent of shared variance (%) of BigFLICA decomposition across a range of dimensionalities in the UKB data. Upper triangle: the explained variance of a lower-dimensional decomposition by a higher-dimensional decomposition. Lower triangle: the explained variance of a higher-dimensional decomposition by a lower-dimensional decomposition.

|  | IC25 | IC100 | IC250 | IC500 | IC750 |
| --- | --- | --- | --- | --- | --- |
| IC25 | 100.00 | 99.98 | 99.99 | 99.99 | 99.99 |
| IC100 | 88.57 | 100.00 | 99.97 | 99.98 | 99.99 |
| IC250 | 86.38 | 96.99 | 100.00 | 99.98 | 99.99 |
| IC500 | 87.77 | 95.45 | 96.79 | 100.00 | 99.97 |
| IC750 | 85.33 | 95.31 | 96.91 | 99.65 | 100.00 |

**Table S5:** A description of 47 Modalities of UKB dataset used in this paper.

| Abbreviation | full description |
| --- | --- |
| rest k (k=1-25) | Dual regression between IC k of 25 dimensional decomposition of rsfMRI and the whole brain |
| task z1 | Z-statistics of emotion task contrast "shapes" |
| task z2 | Z-statistics of emotion task contrast "face" |
| task z5 | Z-statistics of emotion task contrast "faces>shapes" |
| task c1 | Contrasts of parameter estimate of emotion task contrast "shapes" |
| task c2 | Contrasts of parameter estimate of emotion task contrast "face" |
| task c5 | Contrasts of parameter estimate of emotion task contrast "faces>shapes" |
| TBSS-FA | Tract-Based Spatial Statistics - fractional anisotropy |
| TBSS-MD | Tract-Based Spatial Statistics - mean diffusivity |
| TBSS-MO | Tract-Based Spatial Statistics - tensor mode |
| TBSS-L1 | Tract-Based Spatial Statistics - amount of diffusion along the principal directions 1 |
| TBSS-L2 | Tract-Based Spatial Statistics - amount of diffusion along the principal directions 2 |
| TBSS-L3 | Tract-Based Spatial Statistics - amount of diffusion along the principal directions 3 |
| TBSS-OD | Tract-Based Spatial Statistics - orientation dispersion index |
| TBSS-ICVF | Tract-Based Spatial Statistics - intra-cellular volume fraction |
| TBSS-ISOVF | Tract-Based Spatial Statistics - isotropic or free water volume fraction |
| tracts | summed tractography map of 27 tracts from AutoPtx in FSL |
| VBM | voxel-based morphometry |
| Area | Cortical surface area from Freesurfer |
| Thickness | Cortical surface thickness from Freesurfer |
| Jacobian | Jacobian map of nonlinear registration of T1 image to MNI152 standard space |
| swMRI | T2* image derived from swMRI |
| T2 lesion | White matter hyperintensity map estimated by BIANCA |

